# Coevolution-driven reconstruction of multi-taxa siderophore interaction networks reveals topological diversity of microbial exploitation

**DOI:** 10.64898/2026.07.28.741125

**Authors:** Guanyue Xiong, Ruichen Xu, Yukun Zheng, Linlong Yu, Yufei Qiao, Shizheng Tian, Binglei Wang, Ruolin He, Zihao Yang, Xiaoying Bian, Yang Bai, Huimin Yu, Zhiyuan Li

**Affiliations:** Center for Quantitative Biology, Academy for Advanced Interdisciplinary Studies, Peking University, Beijing 100871, China; Peking-Tsinghua Center for Life Sciences, Academy for Advanced Interdisciplinary Studies, Peking University, Beijing 100871, China; Department of Chemical Engineering, Key Laboratory of Industrial Biocatalysis (MOE), Tsinghua University, Beijing 100084, China; State Key Laboratory for Gene Function and Modulation Research, School of Life Sciences, Peking University, Beijing 100871, China; Peking-Tsinghua Center for Life Sciences, New Cornerstone Science Laboratory, Peking University, Beijing 100871, China; Helmholtz International Lab for Anti-infectives, State Key Laboratory of Microbial Technology, Shandong University, Qingdao 266237, China

**Author notes:** These authors contributed equally to this work (Guanyue Xiong, Ruichen Xu).

## Abstract

Microbial communities are shaped by secreted metabolites that mediate ecological interactions, yet predicting these interactions from genomic sequences remains difficult, because the specific recognition between co-functional metabolites (CFMs), such as siderophores, and their receptor proteins (Rec) cannot be inferred from gene annotation alone. This difficulty arises from three factors: the prevalence of Rec-mediated exploitation, the lack of high-accuracy functional annotations, and the absence of genomic co-localization between functionally paired CFM-Rec in Gram-positive bacteria. Here we present the Coevolution-based Interaction Model (CIM), an automated framework that maps specific CFM-Rec pairings directly from uncurated genomic datasets. Using a dynamic joint optimization strategy that accounts for exploitation asymmetry and avoids combinatorial explosion, CIM identifies functional pairings solely through evolutionary covariation. We validated this approach by reconstructing macroscale iron scavenging networks across nine bacterial taxa. Experiments confirmed that CIM can bridge genomic distances exceeding 3 Mb in the Gram-positive genus *Rhodococcus* to identify unlinked cognate receptors, and can accurately predict cross-utilization by exploiter strains despite substantial receptor sequence heterogeneity in Burkholderiaceae and Rhizobiaceae. Finally, topological analysis of the reconstructed networks shows that siderophore exploitation acts as a universal topological glue, fusing fragmented microbial populations into highly connected communities, and that the exploitability of siderophore production reverses depending on network modularity. CIM thus offers a scalable, sequence-to-ecology approach for predicting interactions mediated by secondary metabolites in microbial communities.

## Introduction

Microbes secrete secondary metabolites into their surrounding microenvironment to shape their niches and enhance their survival[1]. Such self-benefiting functions are typically realized by specific metabolite-protein recognition pairs. A prime example is the siderophore-receptor system: to overcome the scarcity of environmental iron while meeting intracellular demand, microbes synthesize and secrete siderophores, a structurally diverse class of small metabolites that chelate ferric iron in the microenvironment[2, 3]. The resulting siderophore-iron complex is then recognized by specific membrane receptors and imported into the cell, allowing the organism to acquire this essential nutrient. With over a thousand siderophore structures characterized to date[4], these metabolites generally engage their cognate receptors through highly specific “lock-and-key” recognition[5]. Another example is seen in the uptake of vitamin B12, where specialized outer-membrane receptors such as BtuB bind this scarce cofactor with similar high specificity to actively transport it into the cell[6]. Beyond nutrient acquisition, analogous metabolite-protein recognition also underlies quorum sensing, where extracellular autoinducers are detected by cognate sensors to coordinate collective behavior[7]. Collectively, we refer to such extracellular metabolites, whose ecological functions are realized through specific protein recognition, as co-functional metabolites (CFMs), and to their cognate protein partners as recognizers (Recs). Whether manifested as siderophore-receptor, VB12-BtuB, or signaling metabolites-sensor interactions, such CFM-Rec pairs underscore a pervasive biochemical strategy by which microbes interact with their niches.

Because the production and recognition of these metabolites can be genetically separable, CFM-Rec systems exhibit a cost-benefit division, naturally mediating complex social interactions. Usually, metabolite production represents the “cost” side, heavily consuming the cell’s limited protein and energy budgets. For example, although siderophores are relatively small molecules around 500-1,500 Da, most are synthesized by nonribosomal peptide synthetase (NRPS)[8], a diverse class of multi-modular megasynthase assembly lines that easily exceeds 1-2 MDa[9]. Other siderophore are synthesized by NPRS-intendent siderophore (NIS) or hybrid pathways, which also involve multiple enzymes and consume ATPs in each synthetic step[10]. Once secreted, these metabolically expensive CFMs readily diffuse into the microenvironment. Then, the benefit side is claimed by the recognizer (Rec) proteins, which act as the selective harvesting machinery. Iron-siderophore complexes are captured by outer-membrane TonB-dependent receptors (TBDR) in gram-negative bacteria[11] and cell-surface-anchored substrate-binding proteins (SBPs) in gram-positive bacteria [12], vitamin B12 is imported by BtuB, and population information is decoded by quorum-sensing sensors[13].

Crucially, this cost-benefit division drives widespread metabolite exploitation. In addition to encoding “self” Recs for their own CFMs, microbes frequently harbor multiple “exploitative” Recs in their genomes to capture foreign CFMs that they do not synthesize[2, 14, 15]. Coupled with high CFM-Rec specificity and diversity, such exploitation fosters rich ecological dynamics. For example, Strain *A* can outcompete its competitor *B* for iron by secreting an exclusive siderophore that B cannot utilize, and/or exploit B by expressing a foreign receptor dedicated to B’s siderophores[16, 17]. Conversely, mutualistic cooperation arises when both strains produce and share the same siderophore pool, and reciprocal exploitation has been suggested to promote dynamic coexistence [18, 19]. Ultimately, by mapping out these specific CFM-Rec recognition relationships, we gain a powerful blueprint for deciphering microbial interaction networks[5, 20].

With the rapid expansion of microbial genomic data and the advancement of data mining algorithms, we are now approaching the era of predicting CFM-Rec mediated interactions from sequence data alone. Remarkable progress has been made in predicting metabolite structures from biosynthetic enzymes. For NRPS, the megasynthases responsible for synthesizing many complex metabolites including siderophores, early rules deciphering substrate-defining domains[21, 22] have matured into comprehensive pipelines that predict 2D chemical structures directly from sequences[23, 24]. Recently, protein language models (PLMs) like ESM-2 have further advanced this by predicting products directly from 3D binding pocket architectures[25]. Simultaneously, recognizer (Rec) proteins are being systematically mapped across genomes using standardized transporter classifications [26] and specialized pHMM models for iron-uptake systems[27]. Recent genus-focused pipelines have successfully surveyed iron-scavenging machineries across ∼2,000 *Pseudomonas* genomes[14], while function-centered alignment algorithms have yielded a global overview of siderophore synthetase distributions across the bacterial kingdom[28]. Nevertheless, merely identifying “what they produce” and “what receptors they encode” is insufficient; determining the specific lock-and-key recognition between CFMs and Recs remains the critical missing link for reconstructing microbial interaction networks from sequence data alone.

Coevolutionary analysis has emerged as a powerful paradigm for recovering functional interactions from genomic data[29]; however, existing methods often lack the generalizability required to systematically map CFM-Rec networks. Historically, tools like CORASON[30] and genomic co-localization searchers like cblaster[31] have relied on genomic synteny to identify co-functional pairs. While this approach often works in Gram-negative bacteria, in Gram-positive taxa the cognate receptors and their biosynthetic operons are often distant across the chromosome[12]. To overcome this structural limitation, evolutionary co-variation in sequence space provides a more robust metric to link CFMs with their cognate recognizers, building on a fundamental eco-evolutionary assumption: due to the high metabolic cost of CFM synthesis, any producing organism must harbor at least one cognate “self-Rec” to collect the benefit, regardless of how many exploitative receptors it also carries[32, 33]. Driven by cheater-cooperator evolutionary games and host-pathogen arms races, the diversification of CFM-Rec interfaces leaves a detectable co-variation signature across phylogenetic trees. Recently, this co-variation framework was successfully applied to the genus *Pseudomonas*, unveiling an intricate siderophore-mediated interaction network encompassing ∼47 pyoverdine subtypes across nearly 2,000 genomes[34, 35]. However, two major barriers prevent the automated generalization of this framework across the tree of life: first, it currently requires intensive, manual feature annotation of synthetases and receptors[14]; second, the Monte Carlo optimization algorithms used to search for global coevolutionary pairing maxima face severe combinatorial explosion, becoming computationally prohibitive at massive genomic scales.

Here, we present the Coevolution-based Interaction Model (CIM), an automated and generalizable framework designed to reconstruct CFM-Rec interaction networks directly from uncurated genomic data. CIM requires only loosely defined candidate genes or gene clusters as input, tolerating both erroneous entries and imprecise prior classification; instead of relying on any fixed sequence threshold, it jointly optimizes functional-group boundaries and lock-and-key pairings through an iterative, coevolution-guided search. We first benchmarked CIM against the well-characterized *Pseudomonas* pyoverdine system, demonstrating its accuracy and specificity in distinguishing true coevolutionary pairings from background noise.

Applying this pipeline across diverse bacterial taxa, we demonstrated that CIM fundamentally overcomes the limitations of traditional network inference. Experimental validation support CIM’s prediction of long-sought receptor for rhequichelin in *Rhodococcus*, which is genomically decoupled from its biosynthetic cluster by megabases. In Burkholderiaceae and Rhizobiaceae, CIM uncovered a shared schizokinen system with pervasive cross-genus exploitation despite high recepter sequence heterogeneity. Aggregating these validated pairings into nine interspecies iron interaction networks further revealed that exploitation, rather than cooperation, provides the topological glue of microbial iron economies, and that the ecological penalty of siderophore production is set by the modular organization of each taxon’s network. CIM thus converts raw genomic sequences into testable ecological hypotheses, offering a scalable route from sequence data to the interaction architecture of microbial communities.

## Results

### The Coevolution-based Interaction Model (CIM) for automated mapping of CFM-Rec interaction networks

The central task in reconstructing CFM-Rec interaction networks is mapping cross-utilizations between species. Specifically, determining whether a CFM produced by Strain *B* can be recognized and imported by a Rec in Strain *A* (Fig. 1a). Given the millions of microbial genomes sequenced to date, inferring these interactions at the raw individual sequence level is computationally intractable. Fortunately, biological systems exhibit convergence at the metabolite-protein interface: phylogenetically-divergent synthetases frequently assemble identical or structurally analogous metabolites[36] (e.g., enterobactin are synthesized by hundreds of microbial genera[28]), and Recs from distinct taxa frequently share binding specificity despite sequence divergence[11, 17].

**Figure 1.**
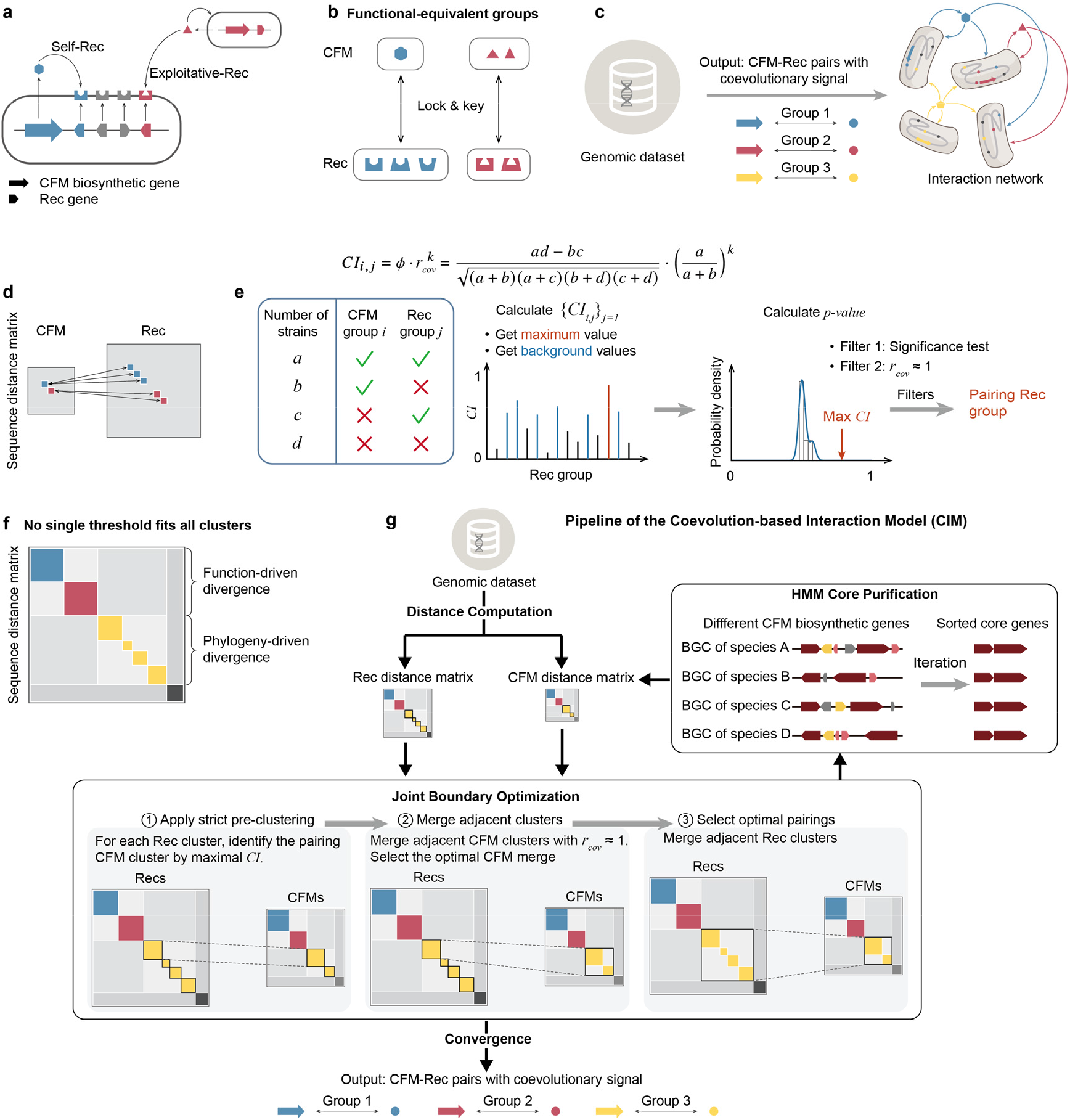
Overview of the Coevolution-based Interaction Model (CIM) for mapping microbial cross-utilization networks. a. Schematic of microbial exploitation strategies mediated by co-functional metabolites (CFMs). Strains encode self-recognizers (Self-Rec) to capture their own CFMs and exploitative-Recs to utilize foreign CFMs. b. Concept of functional-equivalent groups. Sequence-divergent CFMs and Recs sharing equivalent lock-and-key specificities are grouped to simplify network reconstruction. c. The overarching goal of predicting macro-scale cross-utilization networks directly from genomic datasets based on specific CFM-Rec pairings. d. The asymmetry challenge in sequence distance matrices, where a single CFM is targeted by multiple diverse Recs due to pervasive exploitation. e. Formulation of the Coevolution Index (*CI*). *CI* integrates the *φ* correlation coefficient and producer Rec coverage (*r*_cov_) to account for exploitation asymmetry. Candidate pairs must pass dual statistical filters (background significance and near-complete coverage). f. Demonstration of the inherent limitations of fixed-threshold clustering, highlighting how phylogenetic divergence obscure group boundaries. g. The workflow of CIM. The pipeline iteratively performs sequence distance computation, joint boundary optimization (via pre-clustering, boundary merging, and pairing selection), and Hidden Markov Model (HMM) core purification until boundary convergence is achieved, outputting high-confidence CFM-Rec lock-key pairs.

Focusing on this functional convergence, we classify CFM synthetases and Rec proteins into “Functional-Equivalent Groups” (hereafter referred to as CFM and Rec groups) (Fig. 1b). Synthetase and Rec groups are mutually defined by their interaction specificity: all CFMs within a given group exhibit equivalent recognition profiles against all receptors; meaning a Rec that recognizes one member of a CFM group will recognize all members, and vice versa. Consequently, reconstructing the interaction network is streamlined into two interdependent steps: (1) Classification, which partitions synthetase and receptor sequences into discrete functional groups; and (2) Pairing, which establishes specific lock-and-key relationships under the biological constraint of high intra-group specificity and negligible cross-group cross-reactivity. Once these groups and pairs are established, macro-scale cross-utilization networks can be systematically inferred by cataloging the CFM and Rec repertoires encoded within individual genomes (Fig. 1c).

A major challenge in matching CFM-Rec pairs stems from pervasive exploitation. Microbes typically encode far more receptors than synthetases to exploit foreign siderophores[3, 37, 38] (e.g., a single siderophore synthetase versus >10 receptor genes in *Pseudomonas*; Fig. 1d)[14]. This severe numerical asymmetry confounds simple matching. Ideally, if functional groups were perfectly assigned *a* priori, one could pair them simply by maximizing coevolutionary signals. However, accurately defining these group boundaries in isolation is intrinsically difficult. To break this impasse, we developed a joint-optimization approach: using coevolutionary strength itself as the guiding metric to simultaneously define group boundaries and infer their specific lock-and-key pairings.

To drive this joint-optimization, we established a Coevolution Index (*CI*) to quantify the connection between any candidate CFM and Rec group (Fig.1e). Crucially, this index must account for the asymmetry of microbial exploitation: microorganisms synthesizing a metabolically expensive CFM are expected to encode its cognate receptor for self-maintenance, whereas exploitative receptors can occur with or without producing the CFM. To capture this, *CI* is formulated as the product of the *φ* correlation coefficient and the *k*-th power of the Producer Rec Coverage (*r*_cov_), which is defined as the proportion of CFM-producing genomes that also encode the receptor of interest. While this asymmetric metric ensures that true self-utilization pairings yield an *r*_cov_ approaching 1, high *CI* scores can still arise from shared phylogenetic backgrounds rather than authentic functional coupling[39]. Therefore, for a given CFM group, the receptor group exhibiting the maximum *CI* must strictly pass two statistical filters to eliminate false positives: an empirical background evaluation to ensure the signal significantly exceeds phylogenetic noise, and an additional *r*_cov_ threshold to ensure near-complete producer coverage (See SI for detail).

Guided by this filtered *CI* metric, our pipeline addresses the inherent limitation of standard clustering, where no “one-size-fits-all” threshold exists (Fig. 1f). Over-clustering, which splits a functional-equivalent group into multiple clusters, and under-clustering, which merges distinct groups, can both compromise the robust identification of pairing Rec groups (See SI for detail). Moreover, BGCs producing the same metabolites often exhibit various gene organizations, which, combined with imperfect gene annotations, interfere with alignment-based sequence distance measurements by separating BGCs synthesizing the same product into different clusters in sequence space[30]. Therefore, rather than relying on fixed thresholds, our algorithm dynamically infers interactions through a closed-loop heuristic workflow involving the following steps (see Supplementary Methods for algorithmic details):

1. Distance Computation and Pre-clustering: As a prerequisite, pairwise sequence distance matrices are computed for all identified CFMs and candidate Recs. Rec distances are calculated from the specificity-defining feature sequences of TBDR (gram-negative (G-) bacteria) and full-length PBP(specifically PBP2/SBP5) proteins (gram-positive (G+) bacteria), remaining fixed during iterations. Initial CFM sequences are obtained by concatenating all biosynthetic and biosynthetic-additional coding sequences of the BGC; this CFM distance matrix is updated during joint-optimization.
2. Joint Boundary Optimization: Using a strict threshold, these two matrices are fragmented into highly identical pre-clusters with optimal leaf order. To circumvent combinatorial explosion, we employ a boundary-scanning strategy. It progressively expands candidate Rec boundaries at local discontinuities, proposing different candidate Rec groups. Simultaneously, co-occurring CFM groups are proposed, and the Rec-CFM pairs with maximal CI are linked as “candidate lock-key pairs”. Overlapping pairs are resolved by preferring groups with higher compactness and larger sizes.
3. HMM Core Purification: The CFM group of candidate lock-key pairs undergoes an iterative Hidden Markov Model (HMM) purification to extract their uniform core organization of biosynthetic enzymes. If the candidate CFM group successfully converges, it is accepted as a new CFM organization, initiating a new round of CFM distance calculation. If it fails to converge, the CFM-Rec connection is rejected. With the updated CFM distance matrix, Steps 2 and 3 are repeated until convergence.

Through this iterative cycle of candidate generation (Step 2) and CFM reorganization (Step 3), the algorithm repeats the joint-optimization until the boundaries of Rec groups stabilize. Ultimately, this closed-loop pipeline distills raw, uncurated genomic loci into a high-confidence catalog of specifically paired CFM and Rec groups, enabling the systematic reconstruction of macro-scale microbial cross-utilization networks(Fig. 1g).

### Validation of CIM using the established *Pseudomonas* benchmark

To validate our pipeline, we benchmarked the Coevolution Index (*CI*) using the well-characterized genus *Pseudomonas* as a ground-truth dataset. As a Gram-negative bacterium with highly diverse siderophore-scavenging systems, *Pseudomonas* produces a primary siderophore, pyoverdine, which comprises diverse structural subtypes recognized by specific TBDRs termed FpvA. Because the lock-and-key relationships between pyoverdine subtypes and FpvA receptors have been extensively mapped and experimentally verified[14, 34, 35], this genus serves as an ideal reference.

We compiled a curated dataset of 1,182 high-completeness *Pseudomonas* genomes, from which we clustered 1,059 pyoverdine biosynthetic genes into 24 distinct CFM groups (ranked by occurrence frequency, starting from Pyo1; Fig. S1). For recognizers, we collected all 33,573 TBDR sequences across the dataset without relying on prior subtyping, where cognate FpvA receptors accounted for only ∼7.23% of the total pool. This broad definition of Recs was designed to rigorously test our metric’s ability to identify true partners amidst massive background noise.

Using this benchmark dataset, we evaluated the performance of the Coevolution Index (*CI*). Figure 2a and Figure S2 show the biosynthetic gene cluster structures with the corresponding *CI* distribution for these Pyo groups. Among the 24 groups, only two small clusters (Pyo12 with group size 11, Pyo18 with group size 5) failed to rank the previously annotated Rec as the top hit, although the correct Rec still achieved the second-highest score in Pyo12. For three other small groups (Pyo16, Pyo17, and Pyo19), the highest *CI* matched the known cognate Rec, but shared top scores with multiple peaks. This phenomenon arises from phylogenetic autocorrelation in small, highly related populations where self-and non-self-Recs co-occur precisely in the same strains. However, this boundary ambiguity in rare clades does not impair global network reconstruction, as these strains do not serve as sources for outgoing cross-utilization edges.

**Figure 2.**
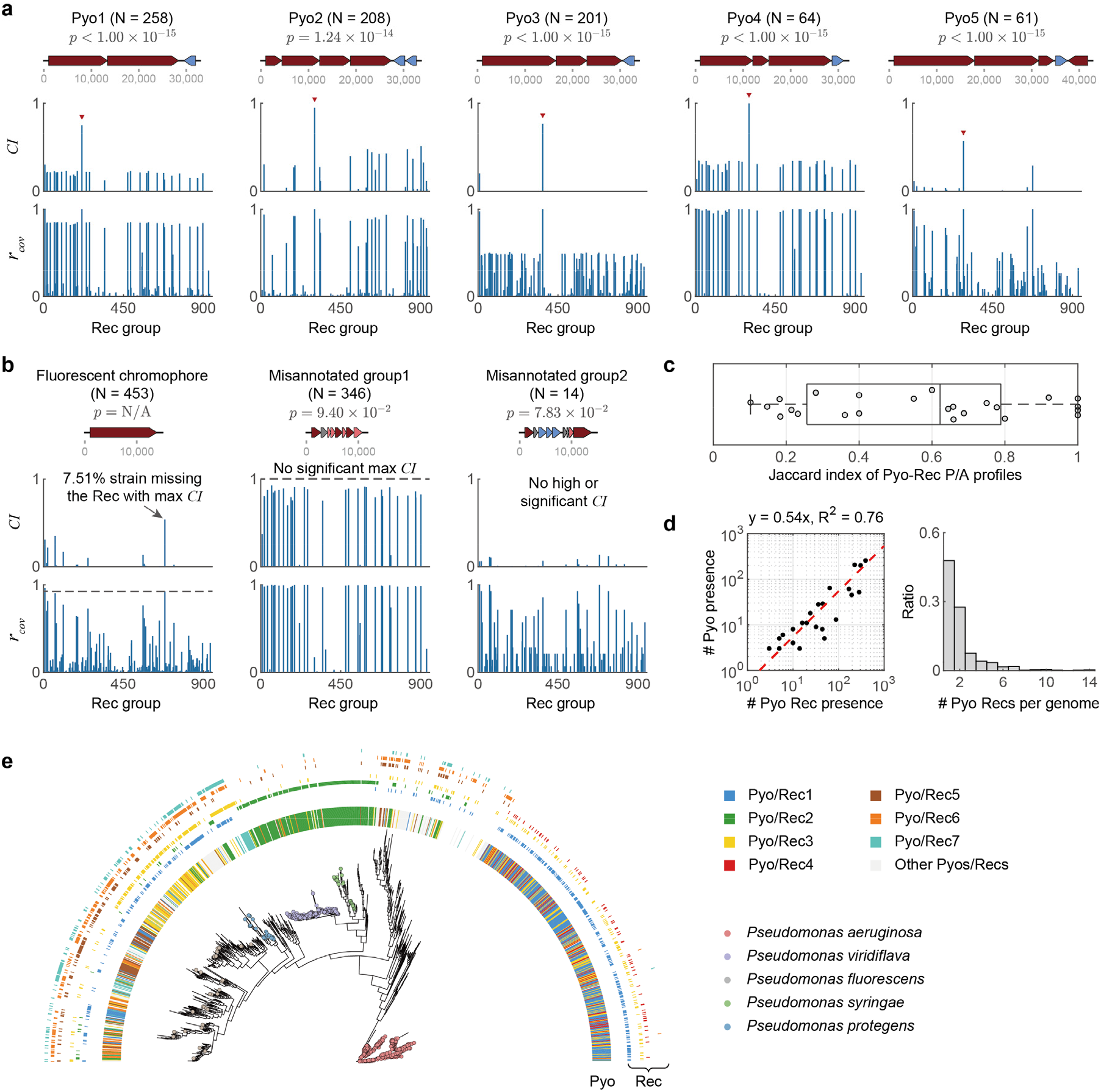
Validation of the Coevolution Index (*CI*) using the *Pseudomonas* pyoverdine benchmark. a. *CI* and *r*_cov_ distributions successfully identify the verified cognate FpvA receptors (red triangles) for major pyoverdine groups (Pyo1–Pyo5). In the gene cluster schematics, dark red arrows indicate core biosynthetic genes and blue arrows indicate receptor genes. b. Evaluation of negative controls, including a conserved fluorescent chromophore and misannotated clusters. These non-interacting groups are effectively rejected by the *CI* statistical filters, demonstrating the metric’s specificity and robustness against genomic noise. c. Jaccard indices of phylogenetic presence/absence profiles for known Pyo-Rec pairs, highlighting the severe degradation of conventional co-occurrence metrics when applied to secondary metabolic systems. d. Quantitative evidence of exploitation asymmetry. The scatter plot (left) reveals that cognate Recs globally outnumber their corresponding Pyos by nearly a factor of two, while the histogram (right) shows a substantial proportion of strains encoding multiple pyoverdine receptors. e. Phylogenetic mapping of major Pyo-Rec pairs across the Pseudomonas tree, illustrating the complex and non-uniform evolutionary distribution of self-and exploitative-receptors.

To further verify the specificity of our metric, we examined the *CI* distribution of three control groups that are misannotated as pyoverdine BGCs. All these groups, not involved in CFM-Rec specificity recognition, displayed patterns distinct from authentic functional pairs (Fig. 2b). For instance, the group representing the fluorescent chromophore (a conserved structural component of pyoverdine not involved in molecular specificity) yielded a false-positive Rec candidate absent in ∼7% of producing strains and was thus rejected by the *CI*’s filter. The other two groups lacked any significant *CI* peaks, confirming that no TBDR exhibits functional coevolution with these groups. Together, these negative controls demonstrate that *CI* effectively filters out genomic noise and avoids false-positive pairings.

We also evaluated whether conventional statistical metrics, such as phylogenetic tree similarity and phylogenetic profiles (PPs) commonly applied in intracellular protein-protein interaction predictions, could reliably identify cognate partners^14,15^. We found that these standard metrics fail in secondary metabolic systems, due to the highly asymmetric distribution of coevolving partners. When we calculated the phylogenetic profiles (PPs) of pyoverdines and their pairing receptors, their Jaccard indices spanned a broad range from 0.1 to 1 (Fig. 2c). This severe degradation occurs because nearly half of the *Pseudomonas* strains (48.39%) encode multiple receptors to exploit foreign pyoverdines they do not synthesize (Fig. 2d, right panel), which introduces massive asymmetric noise that confounds simple co-occurrence analysis. Furthermore, the global abundance of specific Recs is roughly twice that of their cognate Pyos (y = 0.54x, R^2^ = 0.76; Fig. 2d, left panel). Also, the similarity between phylogenetic trees cannot reliably serve as a criterion for identifying pairing Pyo Recs (Fig. S3), due to the shared phylogenetic background of closely related strains. Interestingly, these cheating Recs also do not follow a uniform distribution pattern across the *Pseudomonas* phylogeny. As shown in Figure 2e, Rec1, 3, 5, 6, 7 span a large fraction of clades, whereas Rec2, 4 are more phylogenetically concentrated. In most of the groups, Recs are more random in their phylogenetic spread.

Finally, we evaluated whether our CIM pipeline could reliably resolve functionally insulated clusters from uncurated data without relying on fixed distance thresholds (Fig. S4 and Fig. S5). To test the robustness of the automated “pre-clustering and merging” workflow, we conducted a systematic perturbation test by randomly scrambling the reference cluster boundaries of the *Pseudomonas* dataset. We then applied the CIM procedure to blindly reconstruct the optimal functional partitions from these disrupted inputs. Notably, the dynamically inferred partitions showed strong agreement with the curated ground-truth classifications (Fig. S6). This successful reconstruction demonstrates that CIM overcomes the sensitivity of traditional clustering to arbitrary sequence thresholds, enabling automated, high-confidence mapping of CFM-Rec interaction networks from raw genomes.

### CIM reconstructs siderophore interaction networks across diverse taxa

We applied the CIM pipeline across eight taxonomic levels (one genus *Rhodococcus*, and seven families, including Staphylococcaceae, Bacillaceae, Rhodobacteraceae, Vibrionaceae, Rhizobiaceae, Enterobacteriaceae, and Burkholderiaceae), to systematically predict their siderophore-receptor pairing groups (Fig. 3a). Notably, Burkholderiaceae and Enterobacteriaceae exhibited 10 and 7 potentially novel CFM groups, respectively, that share low similarity with any known siderophore BGCs. We characterized the interaction behavior of individual strains by “siderophore profile”, a digital vector describing the repertoire of siderophores groups they produce and uptake. Further, we summarized the strains’ profiles into three categories of distinct iron-scavenging strategies: dedicated producers (synthesizing and only utilizing their own siderophores), facultative exploiters (producing their own while also exploiting foreign siderophores), and pure exploiters (relying entirely on foreign siderophores without production) (Fig. 3b). In the resulting networks, Rhizobiaceae, Enterobacteriaceae, and Burkholderiaceae showed high diversity in both CFM-Rec groups and siderophore profiles, yielding visually complex iron interaction networks.

**Figure 3.**
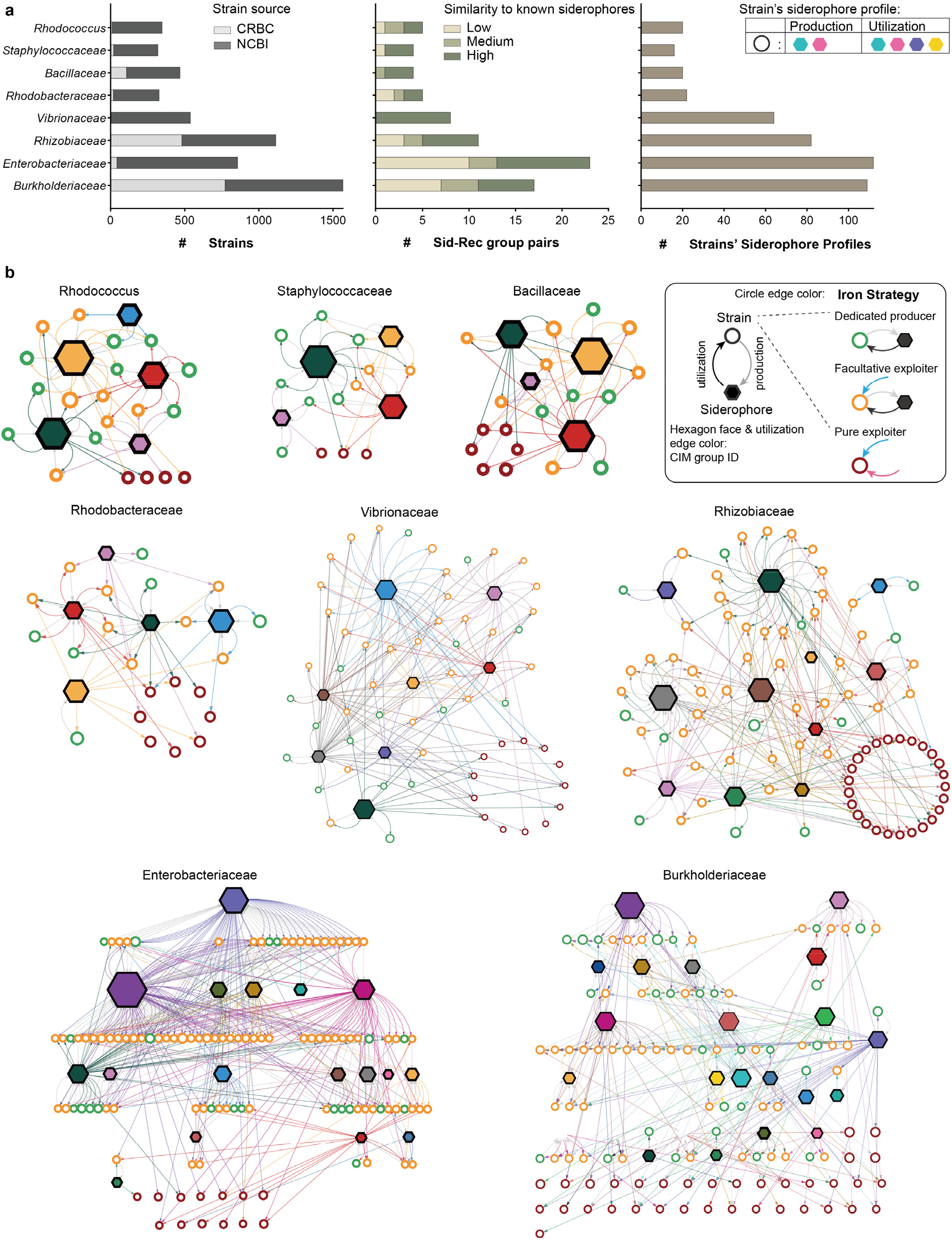
Systematic reconstruction of macro-scale siderophore interaction networks across diverse bacterial taxa. a. Overview of the genomic datasets and CIM-predicted interactions across one genus and seven families. The bar plots quantify the number of analyzed strains (left), the identified CFM-Rec pairing groups (middle), and the overall diversity of strains’ siderophore production and utilization profiles (right). b. Reconstructed iron interaction networks for the eight taxonomic groups. Hexagonal nodes represent functionally equivalent siderophore groups (color-coded by CIM group ID), while circular nodes represent strains’ siderophore profiles. Node sizes are proportional to their prevalence: hexagon size corresponds to the number of producing strains, and circle size reflects the number of strains adopting that profile. Directed edges indicate siderophore production (from circle to hexagon, gray) or receptor-mediated utilization (from hexagon to circle, colored by the CIM group). Strain nodes are highlighted by their distinct iron-scavenging strategies (circle edge color): dedicated producers (green; self-production and utilization), facultative exploiters (yellow; self-production combined with foreign exploitation), and pure exploiters (red; exclusive reliance on foreign siderophores).

We subsequently compared the predicted siderophore-receptor pairs with previously reported systems and found strong consistency. First, in Gram-negative bacteria, although genomic proximity indexes were explicitly excluded from the CIM algorithm, all predicted pairing receptors were located within the siderophore BGCs of the producer strains, except for two low-confidence siderophores in Enterobacteriaceae. Furthermore, our algorithm successfully grouped two previously reported siderophores with similar yet distinct structures (ornibactin and malleobactin) into a single CFM group in Burkholderiaceae, which aligns with previous studies demonstrating reciprocal cross-utilization between their receptors[40, 41] (See Supplement for details). These findings highlight the necessity of classifying CFM-Rec pairs into functionally equivalent groups, indicating that bacterial interactions do not strictly adhere to one-to-one molecular structural correspondence.

In Gram-positive organisms, substrate-binding proteins (SBPs) selectively recognize iron-siderophore complexes, but a substantial proportion of them are not genomically co-localized with their cognate synthetases[12]. For example, the receptors for bacillibactin and schizokinen produced by Bacillaceae are located distant from their respective biosynthetic genes[42, 43]. Notably, our CIM algorithm correctly captured the cognate receptors for both siderophores in Bacillaceae, fully matching the receptor identities and locations confirmed by previous experiments (See Supplement for details).

Finally, to test the broader applicability of our pipeline, we introduced the SBP5 receptor family into the analysis for Bacillaceae and Staphylococcaceae, as it has been reported as a pairing receptor for some opine-like metallophores[44, 45]. We successfully identified the exact pairings for bacillopaline and staphylopine that are consistent with previous experimental reports. This demonstrates that the CIM algorithm is not limited to traditional siderophores but can be broadly extended to other coevolving CFM-Rec systems.

### CIM discovers functional siderophores and links non-adjacent self-recognizers in *Rhodococcus*

Application of CIM to the genus *Rhodococcus* demonstrates the power of our approach in resolving functional pairings where receptors lack detailed curation and reside far from their cognate BGCs. Members of this Gram-positive genus are widely distributed across diverse environments, where siderophores play a vital role in their survival and ecological competitiveness[45, 46]. However, the severe lack of genomic synteny between receptor (Rec) and siderophore biosynthetic genes in Gram-positive bacteria renders proximity-based network inference methods ineffective. Previous characterizations of potential siderophore systems in *Rhodococcus* have remained fragmented and incomplete, leaving a critical disconnect between known biosynthetic genes and isolated chemical structures. Specifically, while two siderophores (heterobactin and rhodochelin) have been fully resolved with both structures and BGCs[47, 48], others lack critical mapping: rhodobactin has an identified chemical structure but an unknown BGC[49], whereas knockout experiments implicate three BGC loci (*iupS*&*iupT* for rhequibactin, *rhbC* for rhequichelin, and *iupU*) in iron-limited survival without corresponding structural characterization of their products[50, 51].

To systematically map the iron-scavenging architectures in *Rhodococcus*, we analyzed a curated dataset of 347 high-quality genomes within this genus, annotating 735 BGC sequences across 20 predicted product classes alongside 8,521 candidate PBP2 receptor sequences. Within two iterative rounds, CIM identified five CFM groups exhibiting significant CFM-Rec coevolutionary signals (Fig. 4a and Fig. S7a-b). Since all five clusters encode NRPS as their core biosynthetic enzymes, we applied PARAS to predict their product structures (Fig. S7b). The phylogenetic distribution of these BGCs across the *Rhodococcus* tree (Fig. 4b) suggests that siderophore biosynthetic capacity is widespread across the genus, with most genomes encoding at least one (30.86%) or two (51.87%) siderophore systems.

**Figure 4.**
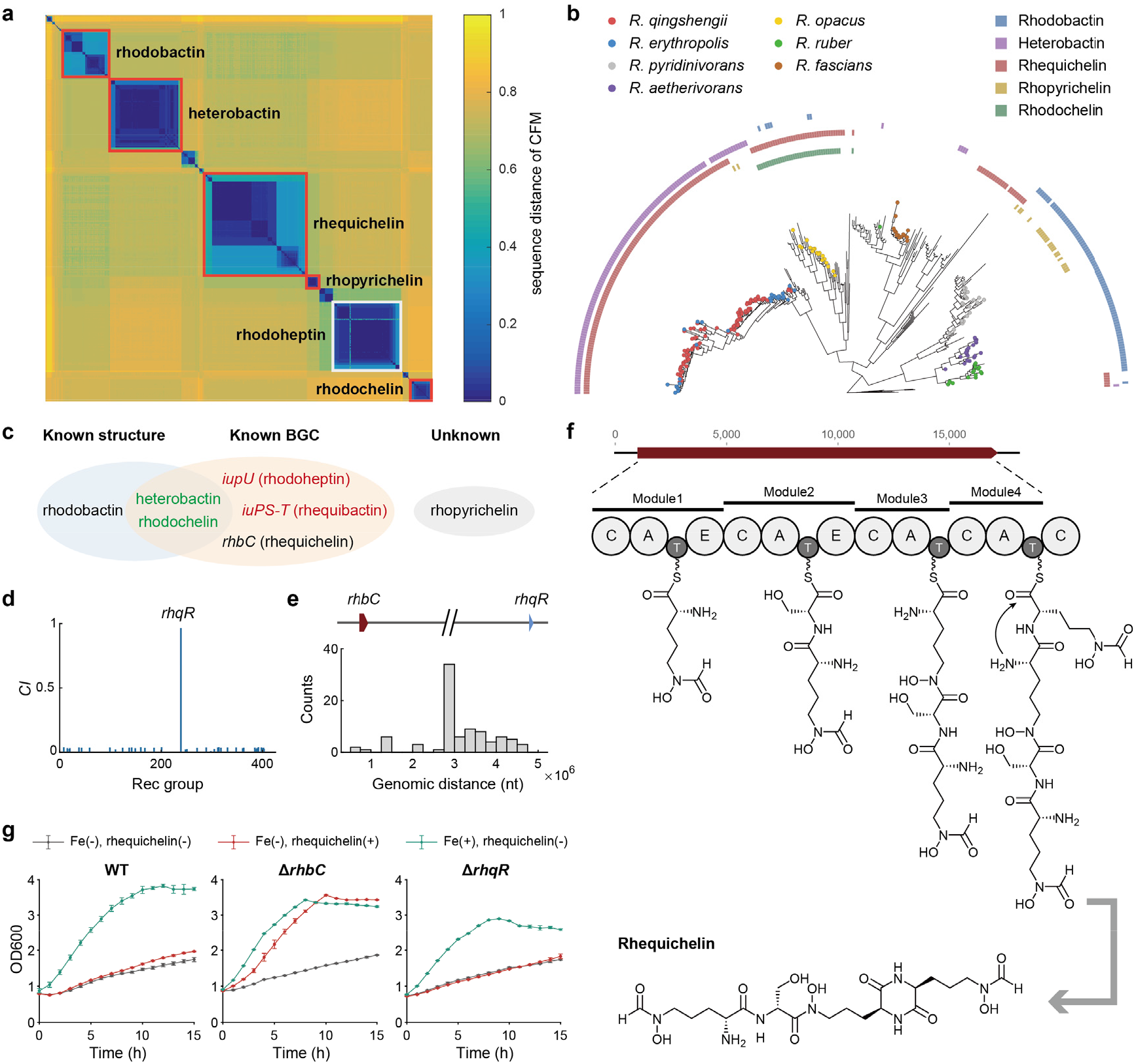
CIM bridges massive genomic distances to identify unlinked self-recognizers in *Rhodococcus*. a. Heatmap displaying the sequence distance matrix of candidate siderophore BGCs in *Rhodococcus* following CIM optimization. Red boxes outline the five functionally equivalent CFM groups that exhibit significant coevolutionary signals with specific Rec groups. b. Phylogenetic distribution of the five predicted siderophore BGC classes mapped across the *Rhodococcus* tree. c. Reconciliation with existing literature. CIM successfully bridges the disconnect between isolated chemical structures (e.g., rhodobactin) and genetically defined loci (e.g., *rhbC* and *iupS-T*). d. Distribution of *CI* scores for the rhequichelin biosynthetic cluster (*rhbC*), pinpointing *rhqR* as its specific, previously uncharacterized cognate receptor. e. Histogram of physical genomic distances between the *rhbC* and *rhqR* loci across producer strains, highlighting their massive spatial decoupling (averaging >3 Mb). f. The proposed NRPS molecular assembly line and the chemically resolved structure of rhequichelin. g. *In vivo* experimental validation of the CIM-predicted long-range CFM-Rec pair. Growth curves under iron limitation confirm that the *rhbC* cluster is responsible for rhequichelin production (growth defect in Δ*rhbC* is rescued by exogenous supplementation), whereas the genomically detached *rhqR* is physiologically indispensable for its uptake (Δ*rhqR* cannot be rescued).

We systematically matched our five predicted CFM groups against existing literaturess (Fig. 4c). The fully characterized heterobactin and rhodochelin systems were readily identified by our pipeline, with their originally reported BGCs falling within the corresponding CFM groups. For the CFM group associated with rhequibactin (*iupS&T*), CIM confirmed that it significantly coevolve with a paired Rec group. Substrate specificity predictions combined with MS/MS structural fragmentation suggested that this compound is actually a novel variant of rhodobactin (Fig. S8 and Fig. S9), which we designated rhodobactin B (m/z = 441.7, holo-form). These integrated analyses resolve a long-standing literature ambiguity, proving that the genetically defined “rhequibactin” cluster is, in fact, responsible for synthesizing rhodobactin-type molecules.

Crucially, CIM demonstrated high accuracy in distinguishing non-siderophore systems by avoiding false-positive predictions. The *iupU* gene locus was previously hypothesized to encode a siderophore due to its role in iron-limited survival. However, CIM detected no PBP2 receptor groups exhibiting significant coevolution with the *iupU*-containing CFM group. This computational rejection is corroborated by structural data: the product of *iupU*, rhodoheptin, lacks canonical iron-chelating moieties (catecholate, phenolate, hydroxamate, or carboxylate; Fig. S10a), and its production is reportedly insensitive to iron concentrations[52]. Together, this supports our CIM prediction that rhodoheptin is unlikely to function as a siderophore. Its necessity for survival under iron limitation may instead stem from participation in a related iron-homeostasis pathway, or its biosynthetic intermediates may regulate other siderophores. The absence of any coevolving PBP2 receptor confirms that CIM effectively filters out physiological noise and avoids misclassifying non-scavenging secondary metabolites (Fig. S10b).

Notably, CIM resolved the long-range genomic decoupling of rhequichelin (*rhbC*) and identified its hitherto unknown cognate receptor, which we termed RhqR (Fig. 4d). In producer genomes, the *rhqR* receptor gene is detached from the *rhbC* biosynthetic cluster, residing at a physical distance averaging over 3,000 kb (Fig. 4e). We isolated and chemically characterized rhequichelin from representative producer strains, resolving its molecular assembly line via MS/MS and adenylation domain profiling (*m/z* = 534.3 [M+H]+; Fig. 4f and Fig. S11). Rhequichelin biosynthesis employs a rare diketopiperazine (DKP) cyclorelease mechanism parallel to erythrochelin (Fig. S12). However, unlike structurally related systems where receptors are locally clustered within the BGCs, the rhequichelin system relies entirely on a genomically unlinked receptor.

To validate that the genomically decoupled RhqR identified by CIM is the true cognate self-recognizer for rhequichelin, we performed both *in vivo* and *in vitro* experiments. Targeted gene knockout in a wild-type producer demonstrated that deleting the biosynthetic cluster (Δ_rhbC_) abolished siderophore production and impaired growth under iron limitation, a defect rescued by exogenous rhequichelin supplementation (Fig. S13 and Fig. 4g), confirming its function as a siderophore. Conversely, deleting the CIM-predicted receptor (Δ_rhqR_) abolished this growth recovery upon supplementation (Fig. 4g), supporting that RhqR is physiologically indispensable for rhequichelin uptake. Complementary *in vitro* biophysical assays confirmed direct, specific ligand binding between purified recombinant RhqR and rhequichelin, establishing a moderate dissociation constant (*K*_d_) of 275.0 nM (Fig. S14). Furthermore, transcriptomic profiling revealed coordinated upregulation of both widely separated loci under iron starvation (Fig. S15). Together, these experimental validations confirm that CIM successfully bridges massive genomic distances to identify authentic CFM-Rec pairs, providing a powerful paradigm for large-scale natural product discovery.

### CIM accurately predicts exploitative cross-utilization despite high receptor sequence heterogeneity

Having validated CIM’s accuracy in identifying cognate pairs within producer genome, we next investigated its ability to predict cross-utilization by “exploiter” strains. To this end, we focused on an uncharacterized BGC (lacking a homologous counterpart in the antiSMASH database) identified in both Burkholderiaceae and Rhizobiaceae, which exhibited distinct BGC arrangements between the two families (Fig. 5a). Crucially, CIM revealed a high ecological asymmetry: only 43 Burkholderiaceae and 22 Rhizobiaceae strains were predicted to produce this siderophore, whereas 178 and 119 strains, respectively, encoded the predicted exploitative receptor distributed across multiple genera (Fig. 5b-c). We isolated two strains whose genomes solely encoded this specific siderophore BGC and purified the active products from culture supernatants under iron-limited conditions. MS/MS analysis and NMR spectroscopy confirmed that the structure was identical to the known siderophore schizokinen (Fig. 5d). While the BGC for this siderophore had previously been reported only in *Anabaena* and *Streptomyces*, this study marks the first report of its BGC architecture in Burkholderiaceae and Rhizobiaceae, revealing an organization distinct from previously described systems.

**Figure 5.**
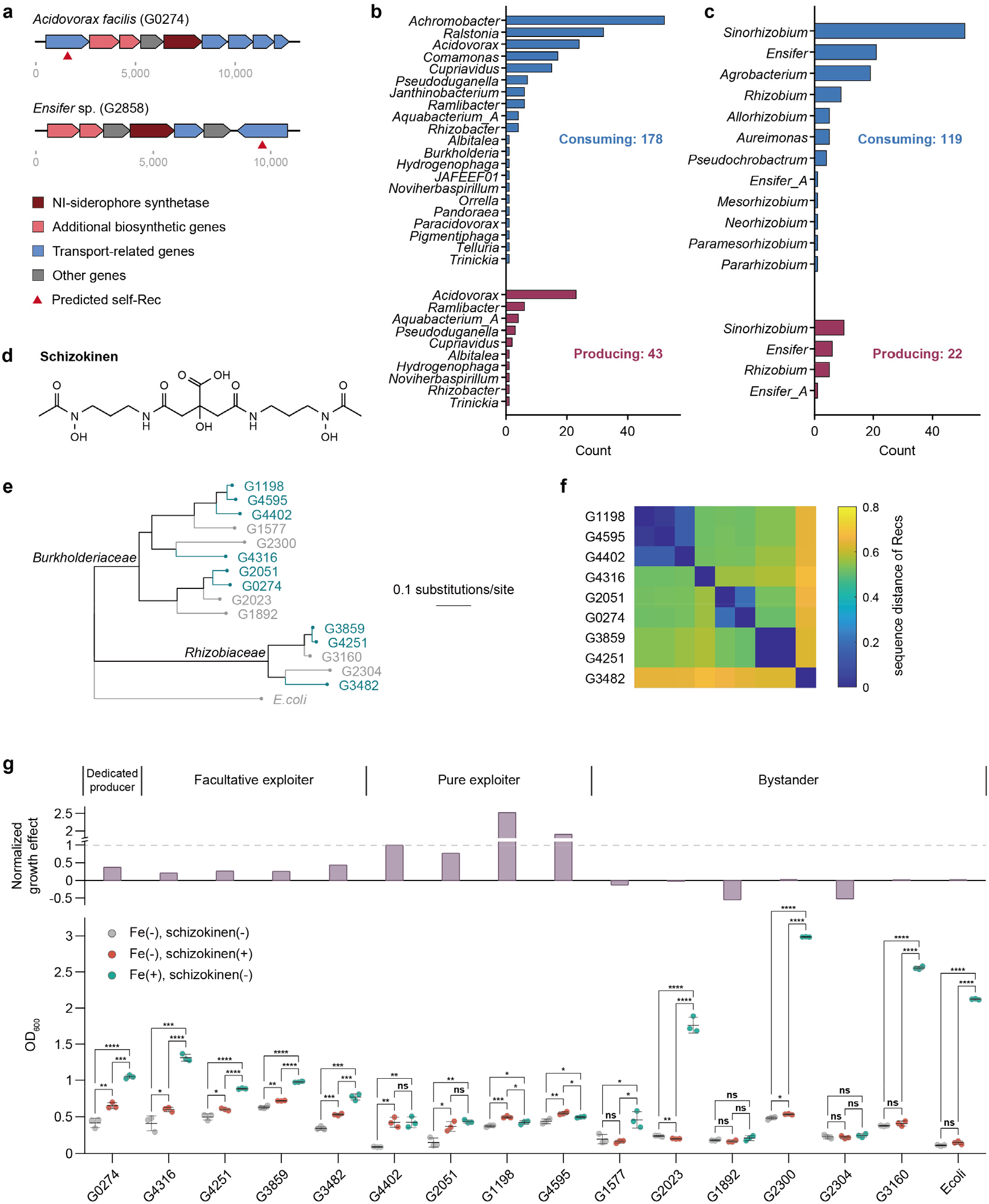
CIM accurately predicts exploitative cross-utilization of schizokinen despite severe receptor sequence heterogeneity. a. Distinct genomic organizations of the schizokinen biosynthetic gene cluster (BGC) identified in representative strains of Burkholderiaceae (*Acidovorax facilis*) and Rhizobiaceae (*Ensifer* sp.). b. Taxonomic distribution of predicted schizokinen producers (red) and exploiters (blue) across genera within the families Burkholderiaceae. c. Same as (b), just for Rhizobiaceae d. Chemical structure of schizokinen, as confirmed by MS/MS and NMR spectroscopy from the isolated culture supernatants. e. Phylogenetic tree of the 15 selected Burkholderiaceae and Rhizobiaceae strains, alongside an *E. coli* control, used for experimental validation. f. Sequence distance matrix of the full-length schizokinen receptors from the predicted utilizing strains, illustrating extreme sequence heterogeneity that confounds traditional homology-based predictions. g. Experimental validation of CIM-predicted cross-utilization via feeding assays. The bottom panel shows absolute growth (OD600) under iron-limited (grey), schizokinen-supplemented (red), and iron-replete (green) conditions. The top bar plot displays the normalized growth effect of schizokinen supplementation. Strains are grouped by their distinct ecological iron-scavenging strategies: dedicated producers, facultative exploiters, pure exploiters, and non-utilizing bystanders.

To experimentally validate these predictions, we randomly selected 10 Burkholderiaceae and 5 Rhizobiaceae strains from the crop root bacterial genome collection (CRBC). This panel included predicted producers, facultative exploiters, pure exploiters, non-utilizing bystanders, and an *E. coli* strain serving as a distantly related control (Fig. 5e). Notably, we observed that schizokinen utilization does not strictly adhere to species boundaries. For instance, within *Sinorhizobium meliloti_A*, strains G3859 and G4251 were predicted as exploiters, whereas G3160 was classified as a non-utilizing bystander. Constructing a sequence distance matrix for the full-length schizokinen receptor genes across these strains revealed significant heterogeneity(Fig. 5f). At a sequence *p*-distance threshold of 0.5, the sequences clustered into five distinct groups, with the receptor of strain G3482 exhibiting an average divergence of 0.65 relative to the others. Such degree of sequence heterogeneity renders traditional homology-based methods incapable of predicting whether these divergent genes perform equivalent functions.

We subsequently purified schizokinen from strain G0274 and conducted cross-feeding experiments on these 16 strains (Fig. 5g). Strains capable of utilizing schizokinen but not producing it were further categorized into “facultative exploiters” (capable of producing other alternative siderophores) and “pure exploiters” (lacking any genomic evidence of siderophore production). As predicted, both producers and exploiters experienced significant growth promotion upon the addition of exogenous schizokinen. In contrast, the bystander group exhibited little to no growth promotion, or even suffered growth inhibition due to the exogenous siderophore sequestering trace iron from the medium. Notably, pure exploiters displayed growth promotion comparable to, or even greater than, the dedicated producers, whereas facultative exploiters exhibited a more moderate response. This pattern may reflect two ecological strategies, where facultative exploiters secure their niches independently by synthesizing proprietary siderophores, whereas pure exploiters expand rapidly once a compatible public good becomes available. Ultimately, these results demonstrate that CIM-predicted interactions are highly reliable, and that integrating coevolutionary signals can uncover complex, asymmetrical cross-utilization networks that are fundamentally invisible to producer-centric annotations.

### Exploitation connects siderophore interaction networks and reflects diverse ecological strategies

Validated by experimental evidence, the siderophore interaction networks predicted by CIM demonstrate high reliability. This resolution allows us to investigate the commonality and specificity of iron-scavenging network architectures across various taxa.

We first evaluated the pervasiveness of exploitation among different siderophores across the nine taxa (Fig. 6a). When plotting production frequency against utilization frequency, many siderophores fall into the region below the diagonal (lower-right), highlighting the widespread existence of exploitative strains capable of utilizing a siderophore without contributing to its production. To quantify this trend, we defined “siderophore exploitability” as the fraction of exploitative receptors among all receptors targeting a specific siderophore (ranging from 0 to 1). Cross-taxa, this index averages 0.34, indicating a widespread yet balanced presence of exploitation. Certain prevalent siderophores are strictly privatized and rarely exploited. For instance, staphyloferrin A in Staphylococcaceae is synthesized by 84.5% of the strains but is almost never pirated, and rhequichelin in *Rhodococcus* is synthesized by 63.0% of the genomes with a low exploitability of 0.03. Conversely, many siderophores known to be crucial for microbial functions exhibit high exploitability. Notable examples include schizokinen (exploitability 0.53 in Bacillaceae, 0.82 in Rhizobiaceae, and 0.76 in Burkholderiaceae), desferrioxamine E in Enterobacteriaceae (0.66), and staphyloferrin B in Burkholderiaceae (0.74). Notably, the exploitability of a siderophore shows no significant correlation with its production pervasiveness, utilization pervasiveness, or synthetic pathway type.

**Figure 6.**
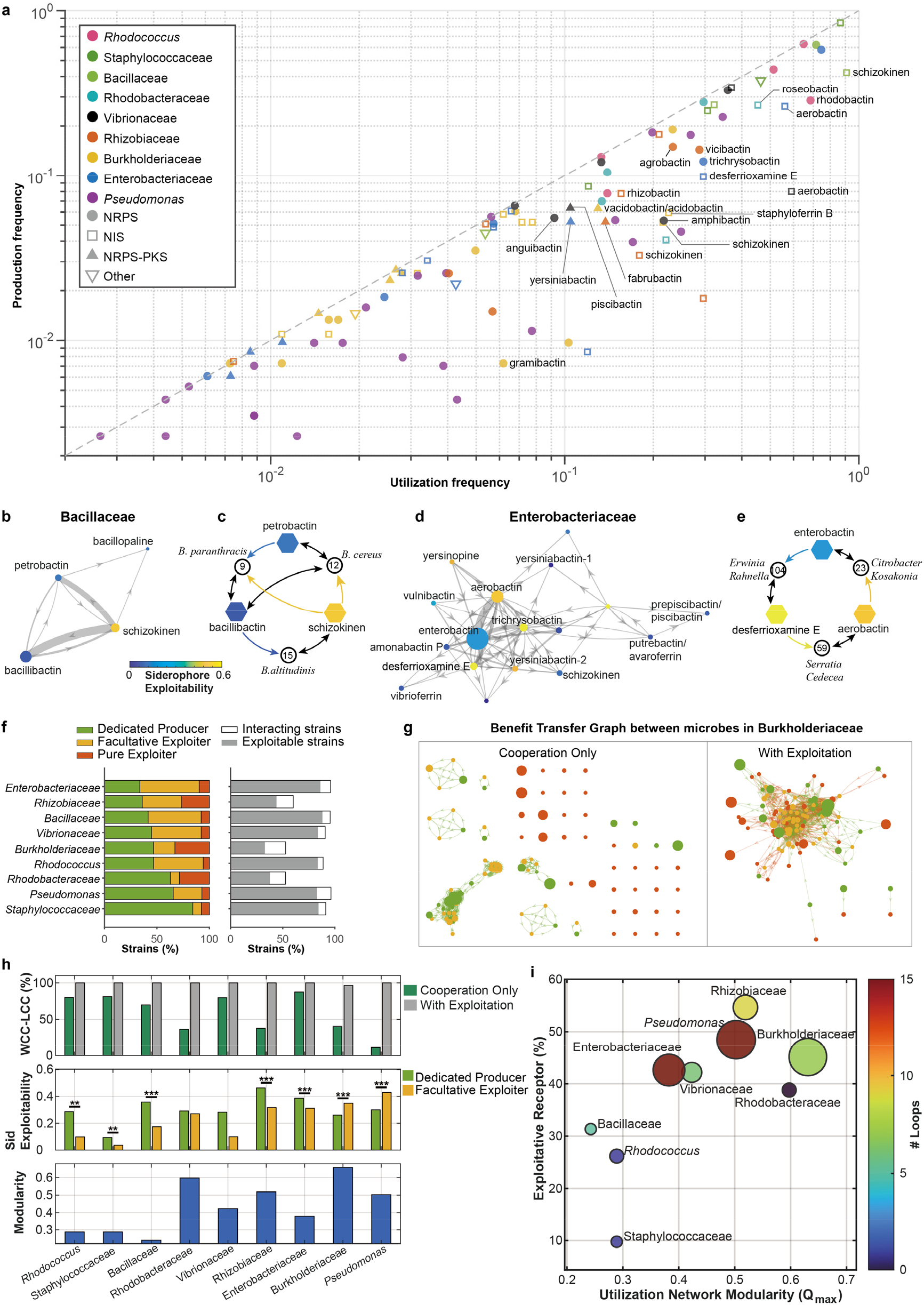
Exploitation connects siderophore interaction networks and reflects distinct ecological strategies. a. Scatter plot comparing the production and utilization frequencies of siderophores across the nine bacterial taxa. Siderophores falling below the diagonal exhibit high exploitability, meaning they are frequently utilized by non-producing strains. b. Siderophore-level Benefit Transfer Graph (BTG) illustrating exploitation among distinct siderophores in Bacillaceae. Circular nodes represent siderophores, with colors indicating their exploitability. Directed edges denote benefit transfer (i.e., the producer of the target siderophore exploits the source siderophore). Edge widths are proportional to the occurrence frequency of these interactions at the strain level. c. Representative reciprocal and rock-paper-scissors (RPS) loops extracted from (b). Circular nodes represent distinct strain profiles, and hexagonal nodes represent siderophores (color-coded as in b). Bidirectional black edges denote the production and utilization of a self-siderophore, whereas directed colored edges indicate the exploitation of a foreign siderophore. d. Same as (d), just for Enterobaceriaceae. e. Same as (c), just for Enterobaceriaceae’s example of rock-paper-scissors. f. Distribution of iron-scavenging strategies (left) and the proportion of exploitable strains (right) across taxa, ranked by an increasing percentage of dedicated producers. g. Strain-level BTGs for Burkholderiaceae. Networks constructed exclusively with cooperative edges fracture into isolated islands (left), whereas the inclusion of exploitative edges fuses the community into a massive connected component (right). h. Topological impact and evolutionary penalties of exploitation. Top: Size of the largest weakly connected component (WCC-LCC) showing near-complete connectivity when exploitation is included. Middle: Exploitability risks faced by dedicated producers versus facultative exploiters, which vary significantly depending on network architecture. Bottom: Utilization network modularity (*Q*_max_) across taxa. i. Clustering of the nine taxa based on utilization network modularity (*Q*_max_) and the proportion of exploitative receptors. Bubble colors correspond to the number of network loops, and bubble sizes are proportional to the number of siderophore synthetases.

Exploitation plays a critical role in connecting the overall siderophore networks. To illustrate this, we constructed a siderophore-level Benefit Transfer Graph (BTG) for each taxon[18]. In this network, a directed edge is drawn from siderophore B to siderophore A if a strain producing A exploits B. Notably, almost all siderophore BTGs are fully connected and feature frequent reciprocal loops (Fig. 6b–e). For example, in the relatively simple network of Bacillaceae, reciprocal benefit transfers are mediated by three distinct siderophore profiles belonging to *B. paranthracis*, *B. cereus*, and *B. altitudinis*. Within this community, bacillibactin is produced by *B. paranthracis* and exploited by *B. altitudinis*, while schizokinen is produced by *B. altitudinis* and exploited by *B. paranthracis*, forming a two-node reciprocal loop. Furthermore, petrobactin is produced by *B. cereus* and exploited by *B. paranthracis*, creating a three-node rock-paper-scissors (RPS) loop. The siderophore BTG becomes considerably more complex in Enterobacteriaceae, displaying intensive benefit transfers among well-known siderophores such as enterobactin and aerobactin. A prominent RPS loop in this taxon involves enterobactin, aerobactin, and desferrioxamine E, with the representative strains mediating this interaction distributed across diverse genera including *Erwinia*, *Citrobacter*, and *Serratia* (Fig. 6e).

From the perspective of the microbial community, this moderate level of exploitation functions as a critical “topological glue” that links interaction networks together. As shown in Fig. 6f, the percentage of dedicated producers varies significantly, ranging from approximately 33% in Enterobacteriaceae to 84% in Staphylococcaceae. However, being a dedicated producer does not isolate a strain from siderophore-mediated interactions. In fact, nearly all producers across every taxon are exploitable, meaning that their synthesized siderophores can be utilized by at least one non-producing strain within the same community. To visualize this at the organismal level, we defined a strain-level BTG. In this network, a directed edge from strain *A* to strain *B* exists if strain *B* can utilize a siderophore synthesized by strain *A*. These edges can be either cooperative (both strains produce the same siderophore) or exploitative (strain B utilizes the siderophore but lacks the synthesis genes). Taking Burkholderiaceae as an example, a network constructed solely with cooperative edges fractures into isolated islands; however, when exploitative edges are incorporated, these fragmented islands fuse into a single highly connected component (Fig. 6g). This striking leap in connectivity is a universal feature across all nine taxa. The size of the largest weakly connected component (WCC-LCC) in cooperation-only networks varies. Taxa such as Enterobacteriaceae and Staphylococcaceae exhibit relatively connected cooperative baselines (encompassing 70% to 80% of all strains), whereas *Pseudomonas* and Rhodobacteraceae are highly fragmented (connecting only 20% to 40% of strains). Nevertheless, once exploitative interactions are included, every network achieves near-complete connectivity, with the WCC-LCC encompassing almost 100% of the strains (Fig. 6h, top panel).

Despite the commonality that exploitation unifies the network, the underlying topological architectures of these communities differ substantially, altering the production penalties associated with different siderophore strategies. In taxa with low modularity, such as Bacillaceae and Rhodococcus, the network is highly nested and interconnected. Within these well-mixed communities, dedicated producers suffer a significantly higher siderophore exploitability risk compared to facultative exploiters (Fig. 6h, middle and bottom panels). Conversely, in highly modular networks characterized by niche segregation, such as those of *Pseudomonas* and Burkholderiaceae, this production penalty is reversed. In these compartmentalized networks, dedicated producers are relatively safe from exploitation, whereas facultative exploiters face significantly higher exploitability risks.

Ultimately, these network topologies align with the overarching genetic investments of the taxa. By mapping the proportion of exploitative receptors against network modularity, the nine taxa segregate into distinct macroscopic clusters (Fig. 6i). For instance, Staphylococcaceae maintains an isolated, low-exploitation lifestyle with very few network loops. Enterobacteriaceae forms a dense, low-modularity network with abundant reciprocal loops and a high proportion of exploitative receptors, likely reflecting an intensive and highly shared competitive environment. In contrast, *Pseudomonas* and Burkholderiaceae manage similarly high levels of exploitative receptors and loops but within highly modular, segregated networks. Together, these patterns suggest that the topological characteristics of siderophore interaction networks are deeply intertwined with the overarching lifestyles and ecological niches of these microbial taxa.

## Discussion

For two decades, the dominant computational route to microbial interaction networks has been genome-scale metabolic modeling (GEM), which reconstructs the conserved enzyme networks of primary metabolism to predict exchange fluxes of amino acids, organic acids, and cofactors among community members [53, 54]. This route captures the metabolic baseline of a community: exchange fluxes of amino acids, organic acids and cofactors. Meanwhile, understanding microbial ecosystems also requires mapping how its members compete, exploit, and negotiate via secreted public goods. Here, we introduce the Coevolution-based Interaction Model (CIM) to establish a second route from genome to micro-ecology, grounded in the hyper-variability and coevolution of secondary metabolism. By computing lock-and-key recognition directly from sequences, and reconstructing macro-scale networks of nine diverse taxa, we establish metabolite-receptor recognition as a newly computable, distinct layer of microbial ecology.

Unlike primary metabolism, co-functional metabolites like siderophores are produced at extreme metabolic cost and diversify under two complementary evolutionary forces: Red Queen antagonistic coevolution between producers and exploiters [55, 56], and Black Queen adaptive gene loss, which converts beneficiaries into dependents[57]. This resulting structural hyper-variability encodes game-like social interactions. However, deciphering these interactions via traditional coevolutionary inference, such as iterative pairing algorithms [58] or phylogenetic profiling [59], often fails because these symmetric metrics assume a one-to-one dependency typical of intracellular protein complexes. Public goods break this contract. A siderophore producer must recognize its own product, but an exploiter can stockpile receptors for metabolites it never synthesizes. This exploitation skews the co-occurrence landscape, and tree similarities conflate pairing with shared ancestry. CIM overcomes this by formulating the Coevolution Index around the one dependency that remains obligatory: the coverage of producers by their cognate recognizer. This asymmetry-aware metric converts an ill-posed symmetric matching problem into a well-posed directional one, a paradigm that should become the default for analyzing any public-good system where cost and benefit loci are genetically separable.

Beyond methodological advancements, our cross-taxa comparison demonstrates that network topology dictates the evolution of siderophore exploitation strategies. In well-mixed, low-modularity taxa such as Bacillaceae and Enterobacteriaceae, dedicated producers are the most exposed to exploitation. Conversely, in highly compartmentalized, modular taxa such as *Pseudomonas*, facultative exploiters attempting to cross niche boundaries bear the higher risk. Topology therefore sets the sign of the evolutionary production penalty: modularity shelters production, while mixing subsidizes exploitation. This reversal offers a mechanistic reading of earlier habitat-level observations, which noted that free-living environmental communities form dense iron networks while spatially structured, host-associated ones are small and fragmented[35].

The extremes of this strategy spectrum are equally informative. At one pole, effectively privatized channels like staphyloferrin A in Staphylococcaceae and rhequichelin in *Rhodococcus* show how producers can escape the social dilemma entirely, consistent with theories that privatization stabilizes cooperation[19]. At the opposite pole, rock-paper-scissors exploitation loops, such as enterobactin-aerobactin-desferrioxamine E, realize in natural communities the cyclic benefit transfers that theoretically scaffold stable coexistence[18]. Crucially, exploitation is not purely antagonistic; xenosiderophore uptake can sustain organisms that would otherwise remain unculturable[17]. The exact same network edges that constitute theft among competitors can represent essential subsidies toward dependents, capturing the duality at the heart of the Black Queen dynamic.

Although developed and validated on siderophores, CIM is defined by an ecological constraint rather than a specific protein fold. The algorithm successfully paired ligands with TonB-dependent receptors in Gram-negative taxa, as well as with unrelated substrate-binding proteins (PBP2 and SBP5) in Gram-positive taxa. In principle, any CFM-Rec system featuring a cost-benefit division and a “producer-must-recognize” constraint falls within its scope. Prime candidates for future expansion include vitamin B12 uptake[11], quorum-sensing signal eavesdropping and quenching[7], and bacteriocin-immunity protein arms races[60]. Extending interaction inference to these systems will move the field from a single elemental economy to a broader landscape of chemical negotiation.

To fully realize this expansion, two current limitations must be addressed. The first is taxonomic reach. CIM effectively maps sequence similarity to functional equivalence within a taxon; across family boundaries, deep sequence saturation and strong phylogenetic noise erode the covariation signal. Because nature ignores these boundaries, as demonstrated by schizokinen bridging Burkholderiaceae and Rhizobiaceae, scaling CIM will likely require coupling sequence coevolution to structural evidence via co-folding and pocket-level docking models [25, 61]. The second boundary is experimental throughput. While CIM rapidly generates thousands of predictions, validation via knockout genetics or purified-ligand feeding remains a slow, organism-specific bottleneck. Closing this loop at scale demands high-throughput phenotyping, such as massively parallel droplet microfluidics or metatranscriptomic readouts[62]. We view this prediction-to-validation throughput gap as the primary rate-limiting factor for the field in the coming years.

Metabolic models taught us what microbial communities can exchange; coevolutionary models can now tell us who is allowed to take what. By making metabolite-receptor recognition computable, CIM adds a crucial missing layer between genome and community, the one written in the language of cost, benefit, and deception. As structural prediction and high-throughput phenotyping mature, decoding this secondary metabolism layer will become as routine a starting point for microbiome research as primary metabolic reconstruction is today.

## Acknowledgement

This work was supported by the National Key Research and Development Program of China (No. 2024YFA0919500), Fundamental and Interdisciplinary Disciplines Breakthrough Plan of the Ministry of Education of China (JYB2025XDXM502), National Natural Science Foundation of China (No. T2321001), State Key Laboratory of Microbial Technology Open Projects Fund (Project NO. M2024-05) and LZ was supported in part by the Peking-Tsinghua Center for Life Sciences.

