## Supplementary material for "Coevolution-driven reconstruction of multi-taxa siderophore interaction networks reveals topological diversity of microbial exploitation": SI Appendix

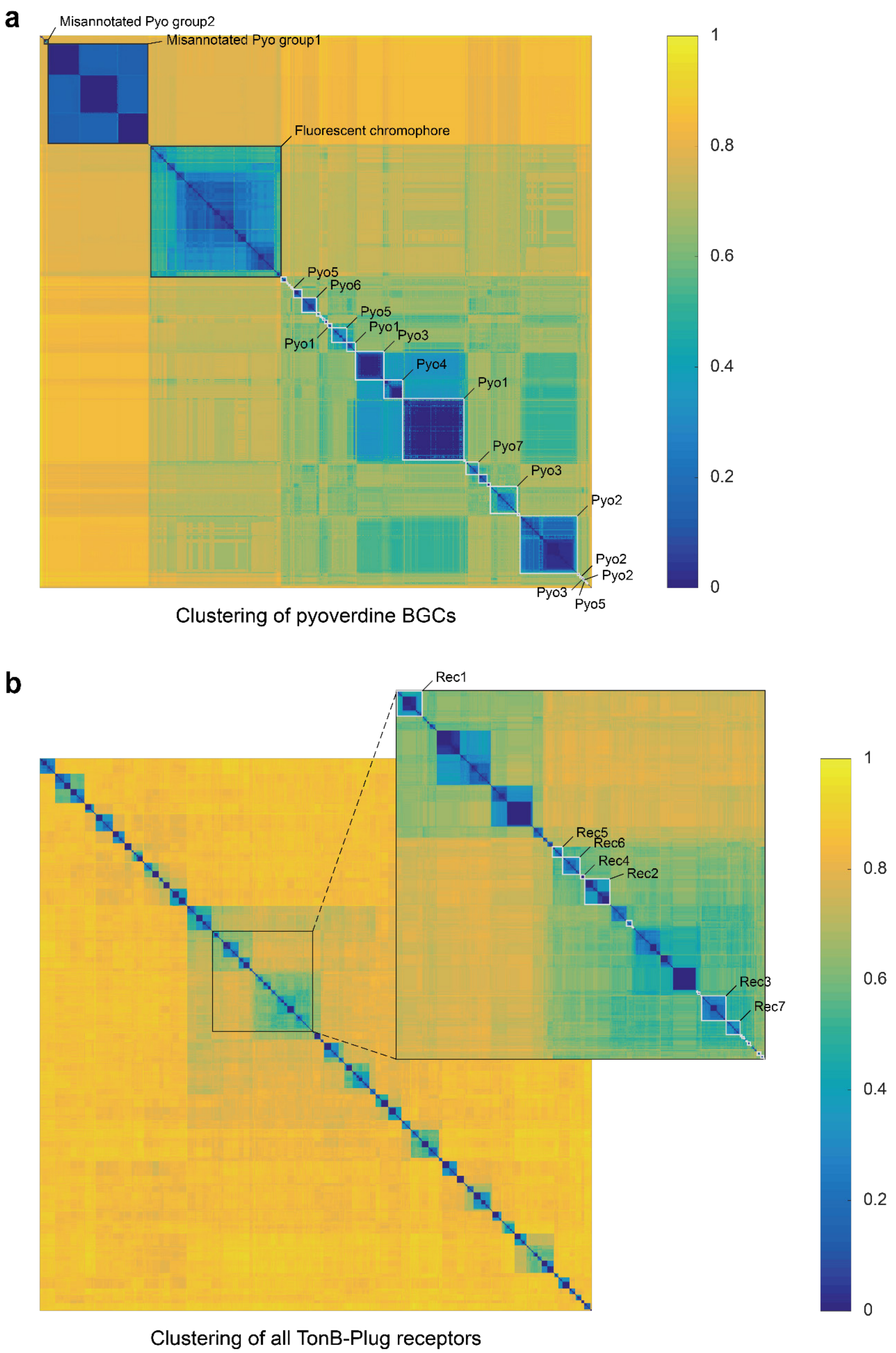


**Fig. S1 | Sequence clustering of pyoverdine systems in *Pseudomonas*. a,** Sequence distance heatmap of pyoverdine biosynthetic gene clusters (BGCs). **b,** Sequence distance heatmap of TonB-dependent receptors, demonstrating the grouping of functionally equivalent modules.

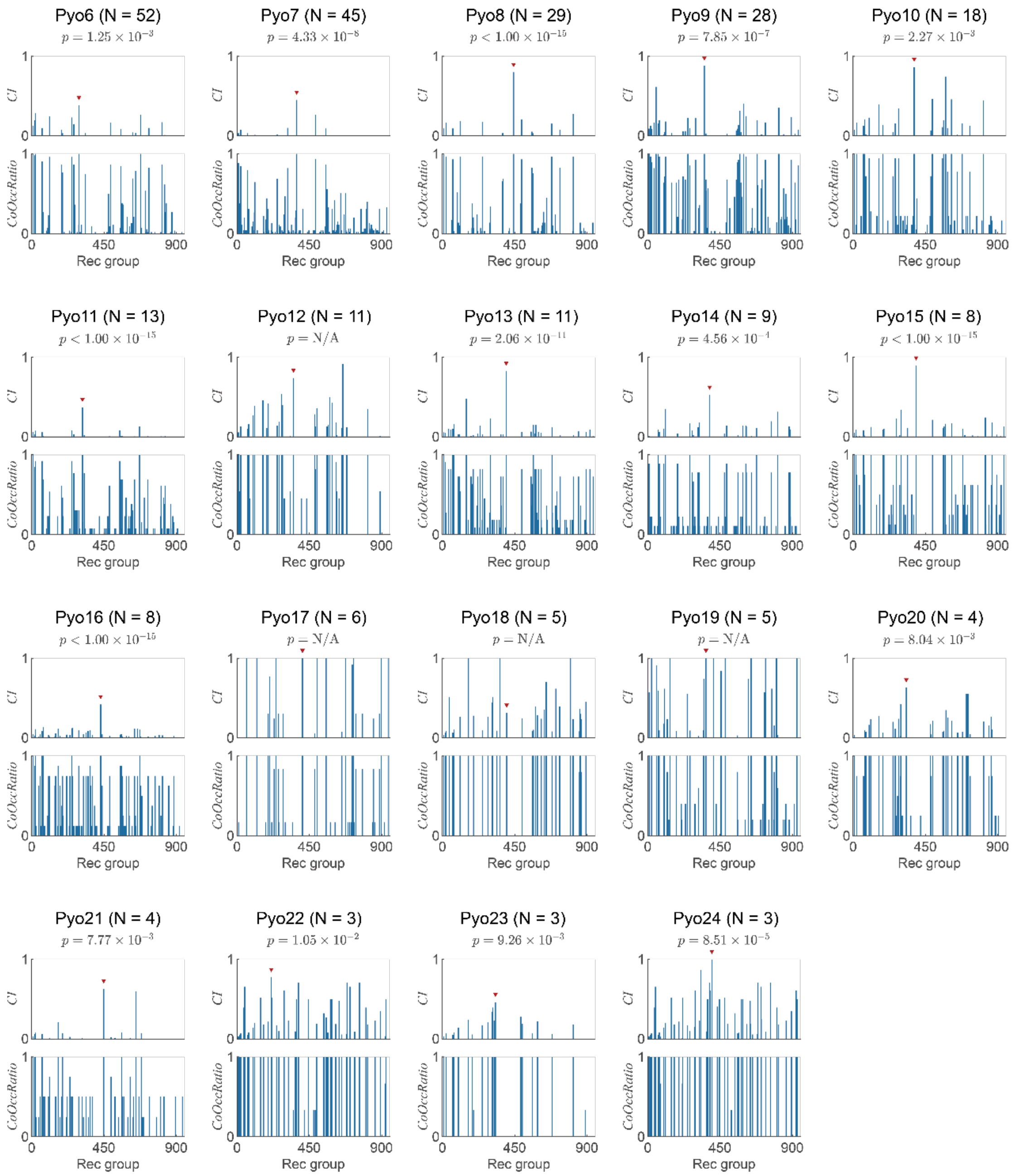


**Fig. S2 | Coevolution Index (CI) distributions across minor pyoverdine groups.** CI and co-occurrence ratio distributions for Pyoverdine groups 6 through 24. Red triangles pinpoint the successfully identified cognate self-receptors.

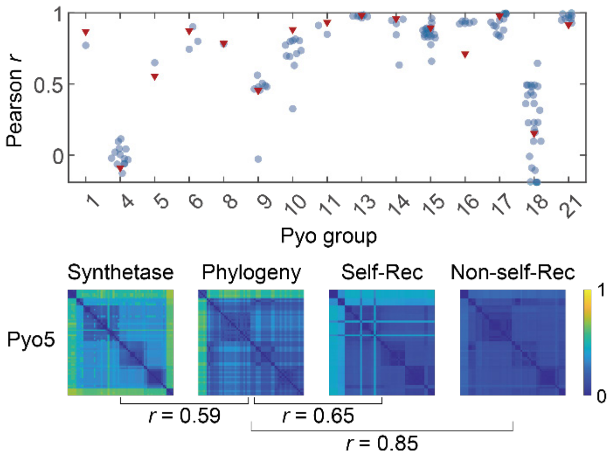


**Fig. S3 | Failure of simple sequence similarity to predict functional pairings.** Pearson correlation analysis demonstrating that overall phylogenetic similarity (Pearson *r*) cannot effectively distinguish true self-receptors from non-self-receptors due to intensive evolutionary noise and horizontal gene transfer.

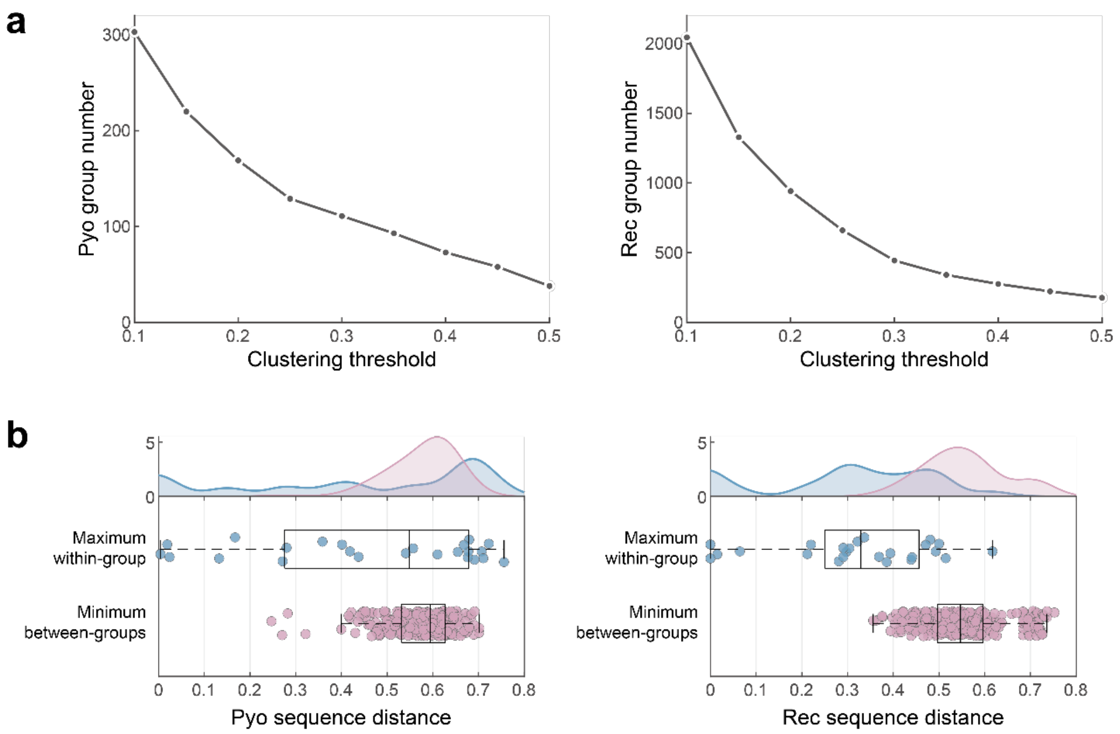


**Fig. S4 | Robustness of the sequence clustering thresholds. a,** The impact of sequence distance clustering thresholds on the total number of generated Pyo and Rec groups. **b,** Density distributions of maximum within-group and minimum between-group sequence distances, confirming clear boundary separations.

**a**

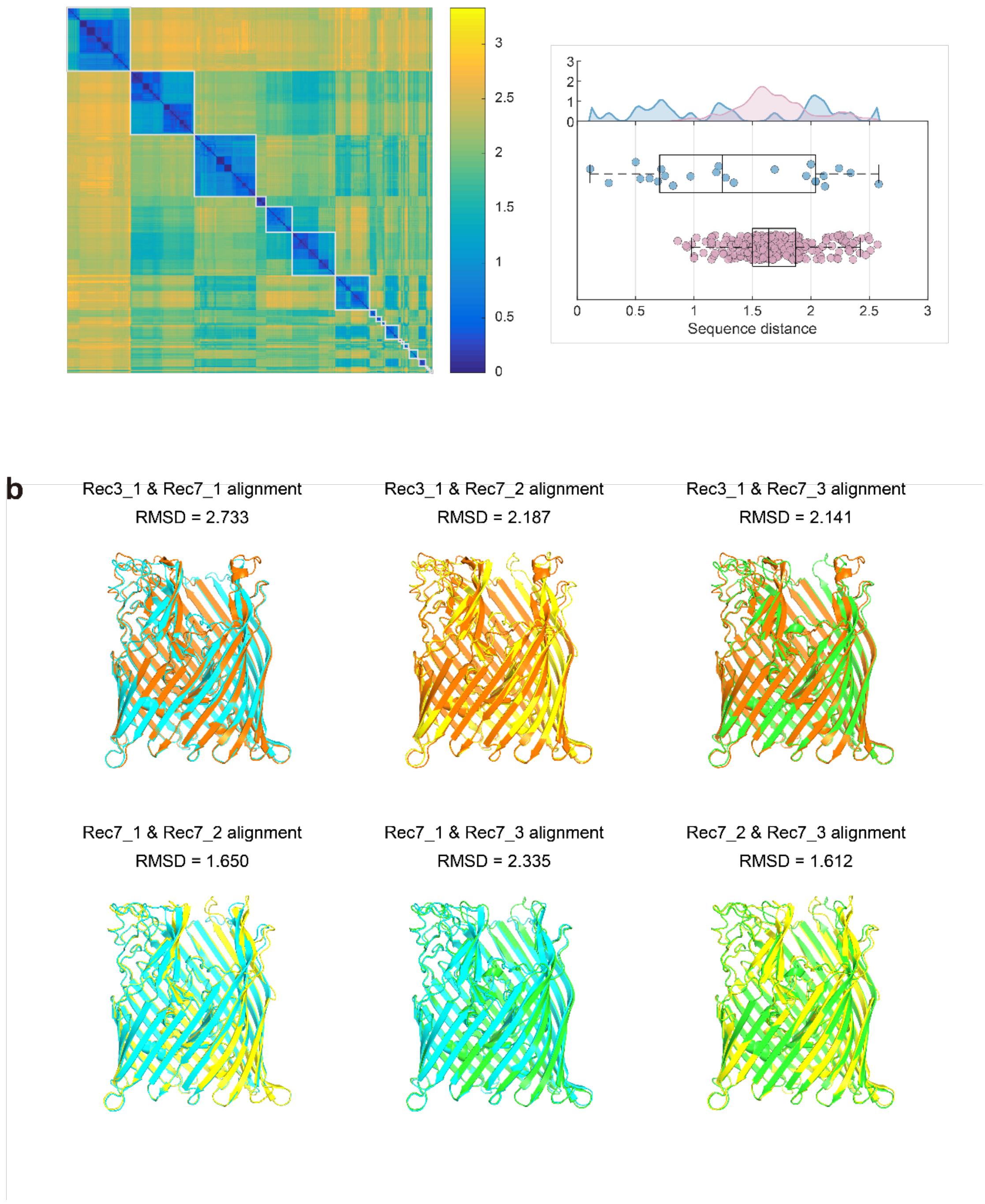


**Fig. S5 | Structural conservation among sequence-divergent receptors.** 3D structural alignments and RMSD values for functionally equivalent but sequence-divergent TonB-dependent receptors (Rec3 and Rec7 groups), illustrating conserved binding pocket architectures despite low sequence identity.

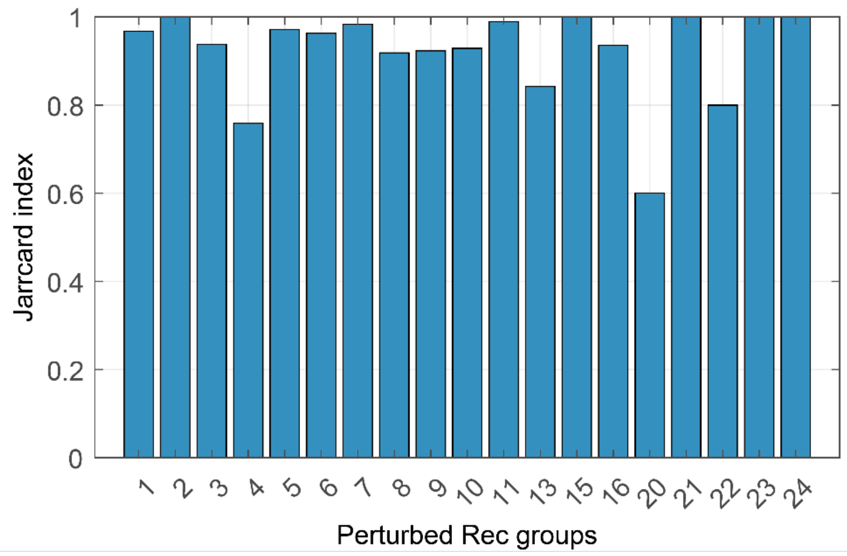


**Fig. S6 | Robustness of CIM predictions to input perturbations.** Jaccard index evaluation showing the high stability of CIM-predicted pairings when subjecting the receptor group assignments to random perturbations.

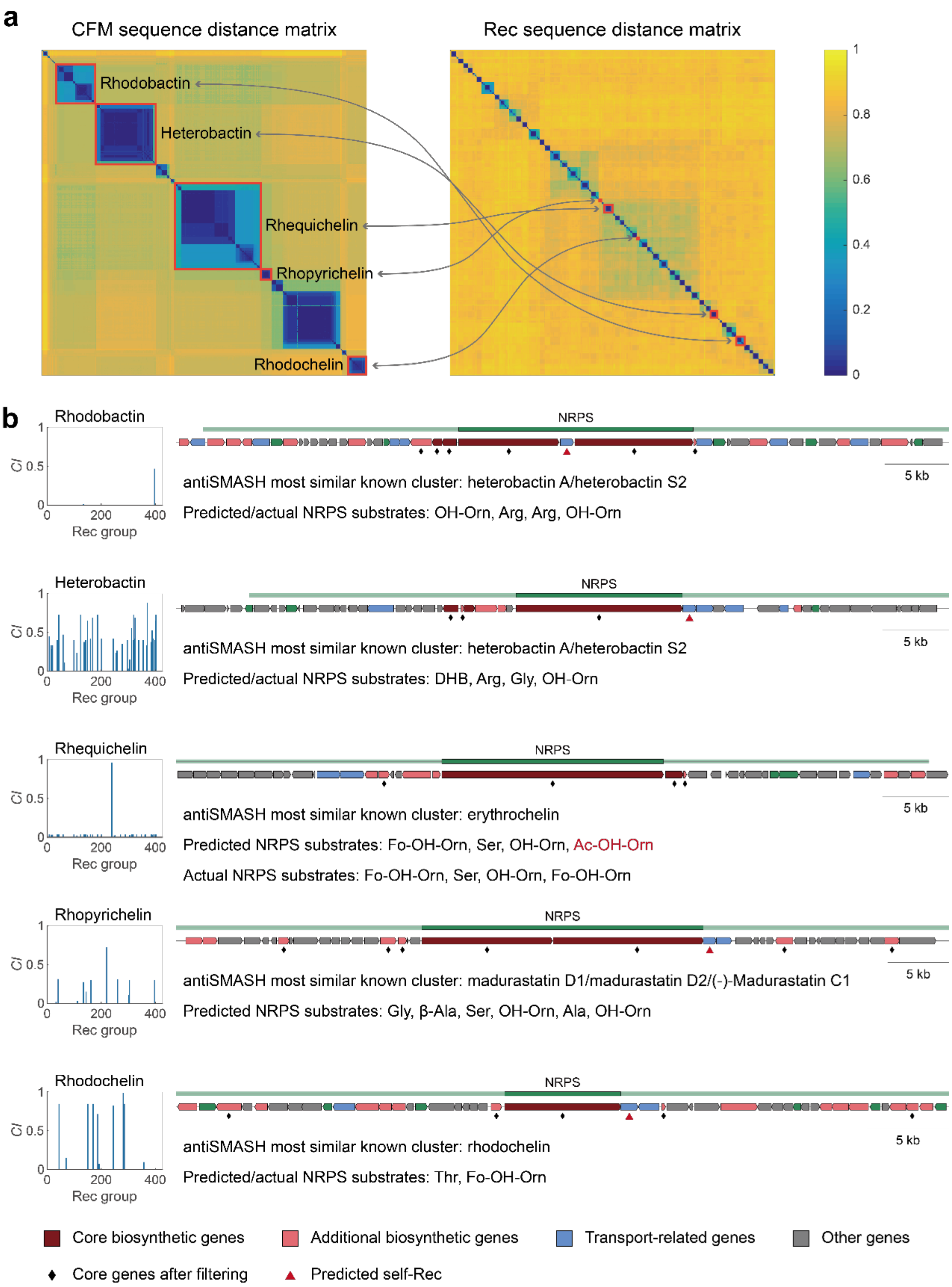


**Fig. S7 | Identification of functionally equivalent CFM-Rec groups in *Rhodococcus*. a,** Sequence distance matrices for candidate siderophore BGCs and receptors in *Rhodococcus*. **b,** Genetic architectures of the five CIM-identified BGCs and their corresponding predicted product substrates.

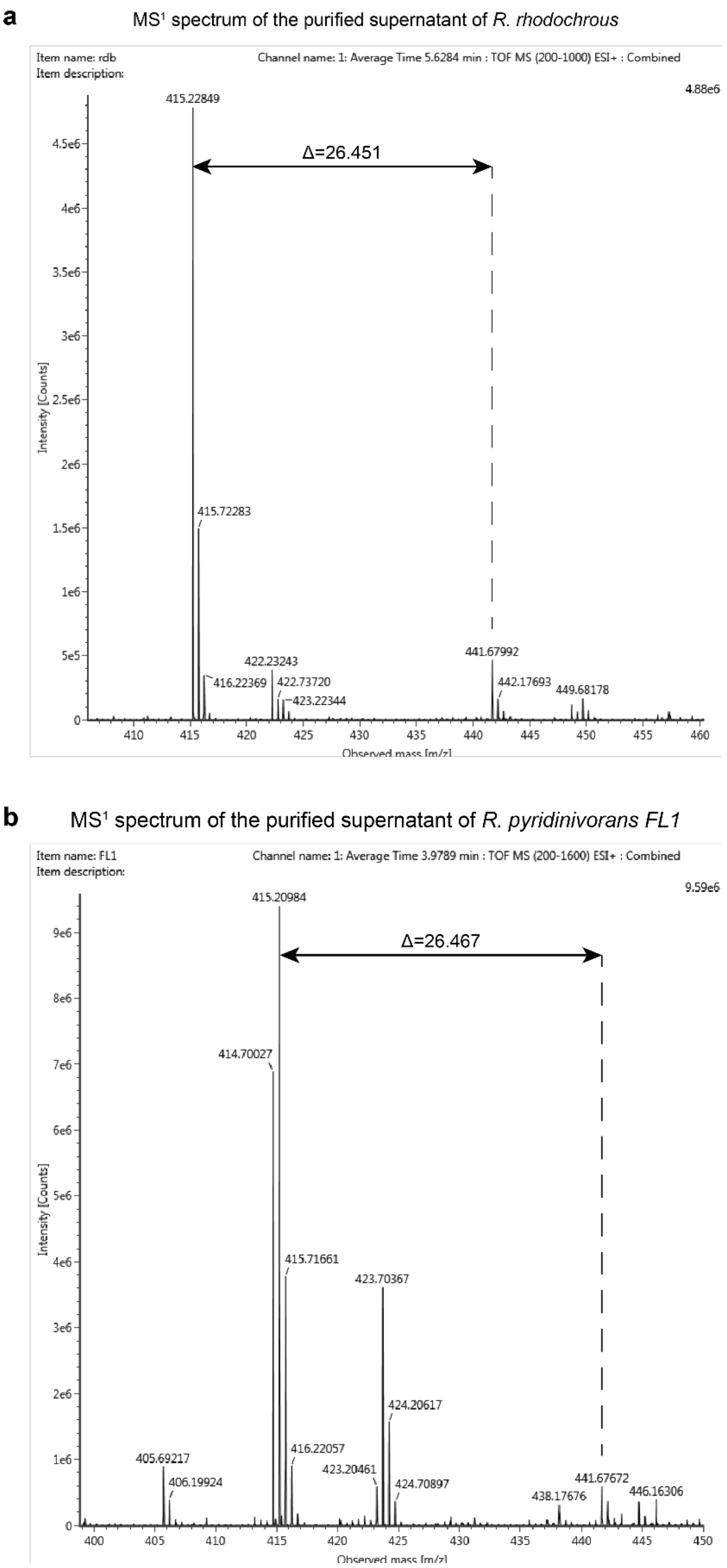


**Fig. S8 | MS1 identification of rhodobactin variants.** MS1 mass spectra of the purified culture supernatants from *Rhodococcus rhodochrous* (**a**) and *R. pyridinivorans* (**b**), identifying the molecular masses corresponding to the rhodobactin family.

a

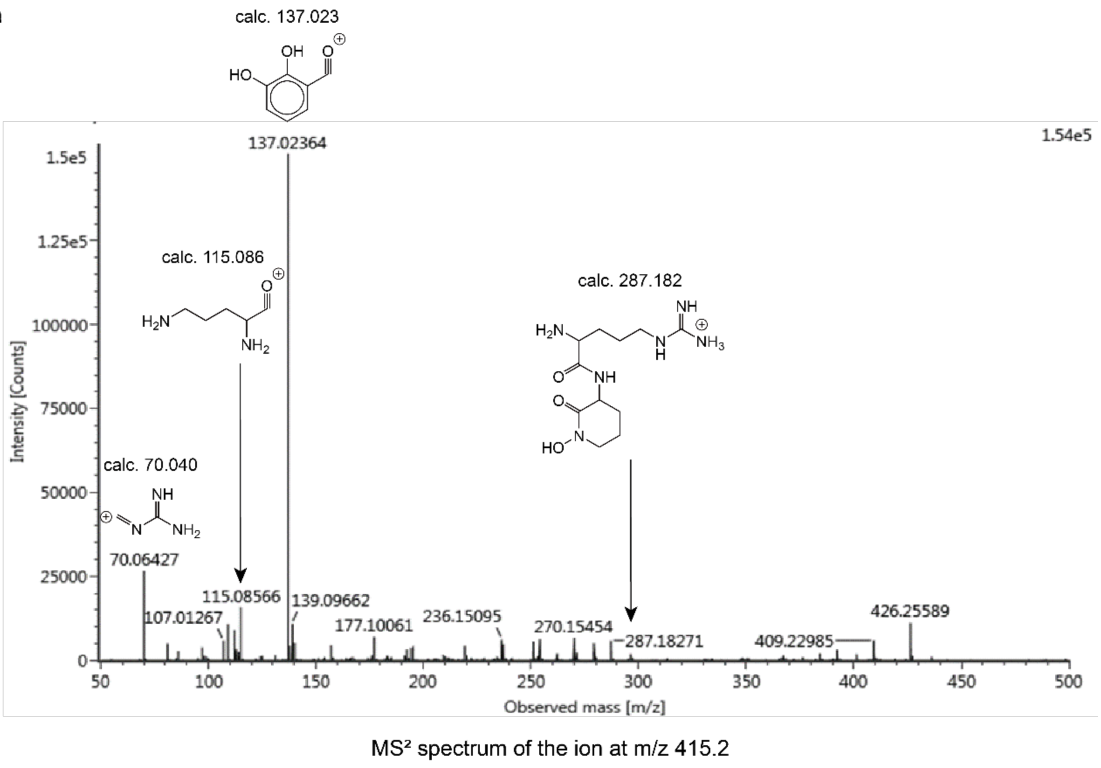


b

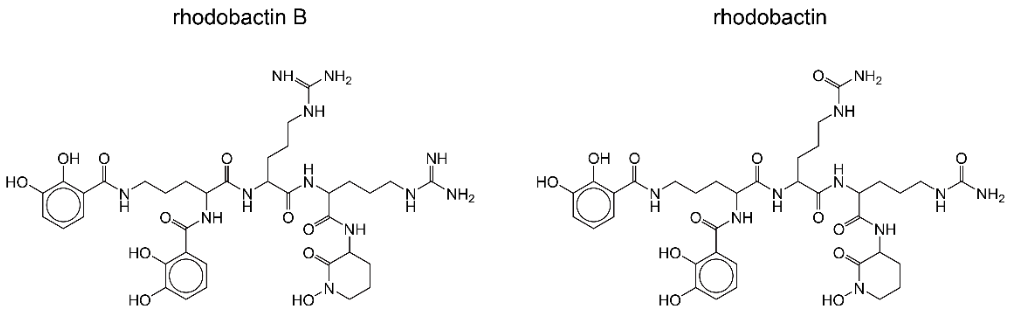


**Fig. S9 | Structural resolution of the novel rhodobactin B. a,** MS/MS fragmentation spectrum of the ion at *m/z* 415.2. **b,** The chemically deduced structure of the novel variant rhodobactin B compared to the canonical rhodobactin.

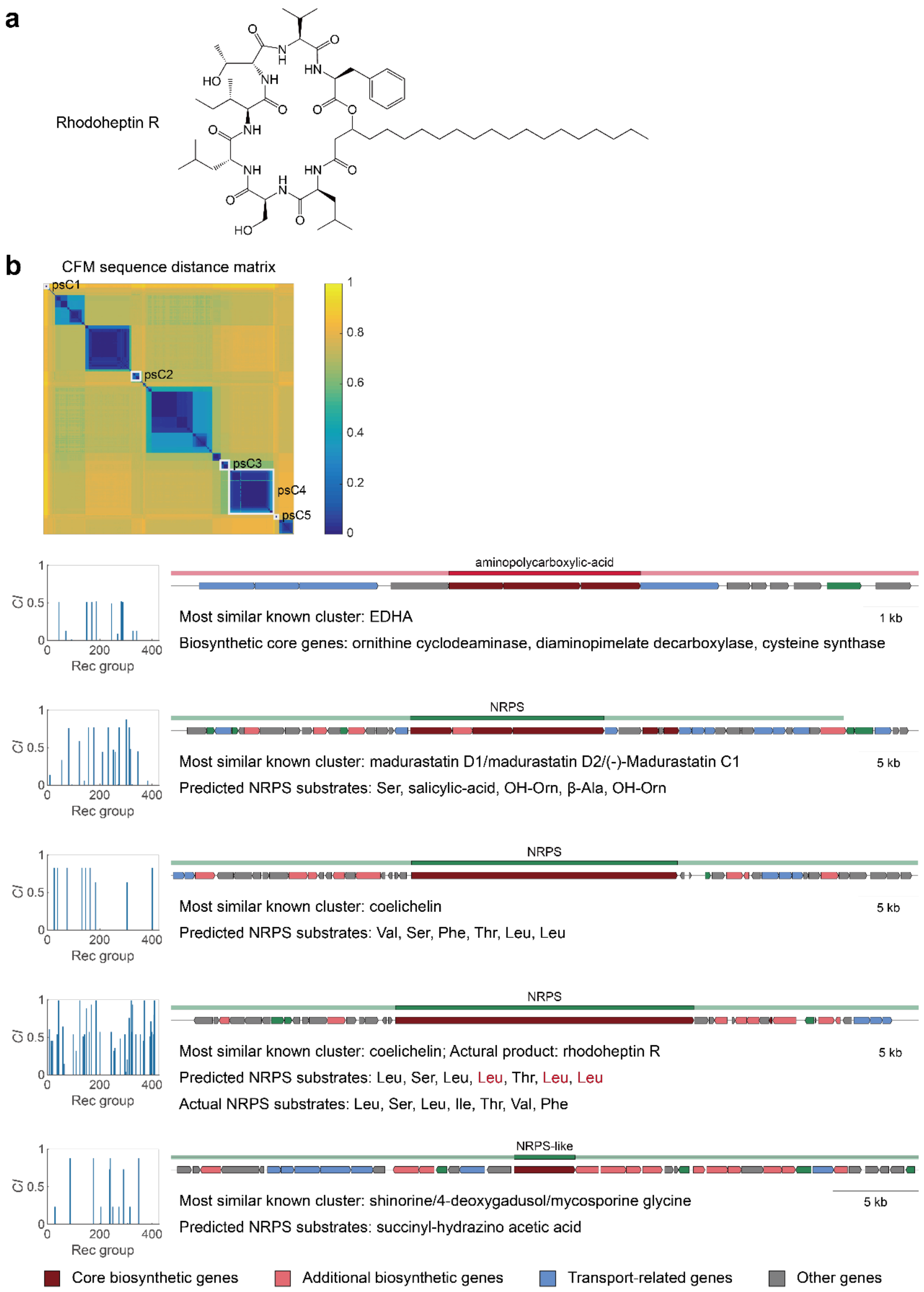


**Fig. S10 | Computational and structural rejection of rhodoheptin as a siderophore.** **a,** Chemical structure of rhodoheptin R, notably lacking any canonical iron-chelating moieties. **b,** CIM sequence distance matrix and CI distributions showing the complete absence of any significantly coevolving self-receptor for the rhodoheptin (*iupU*) cluster.

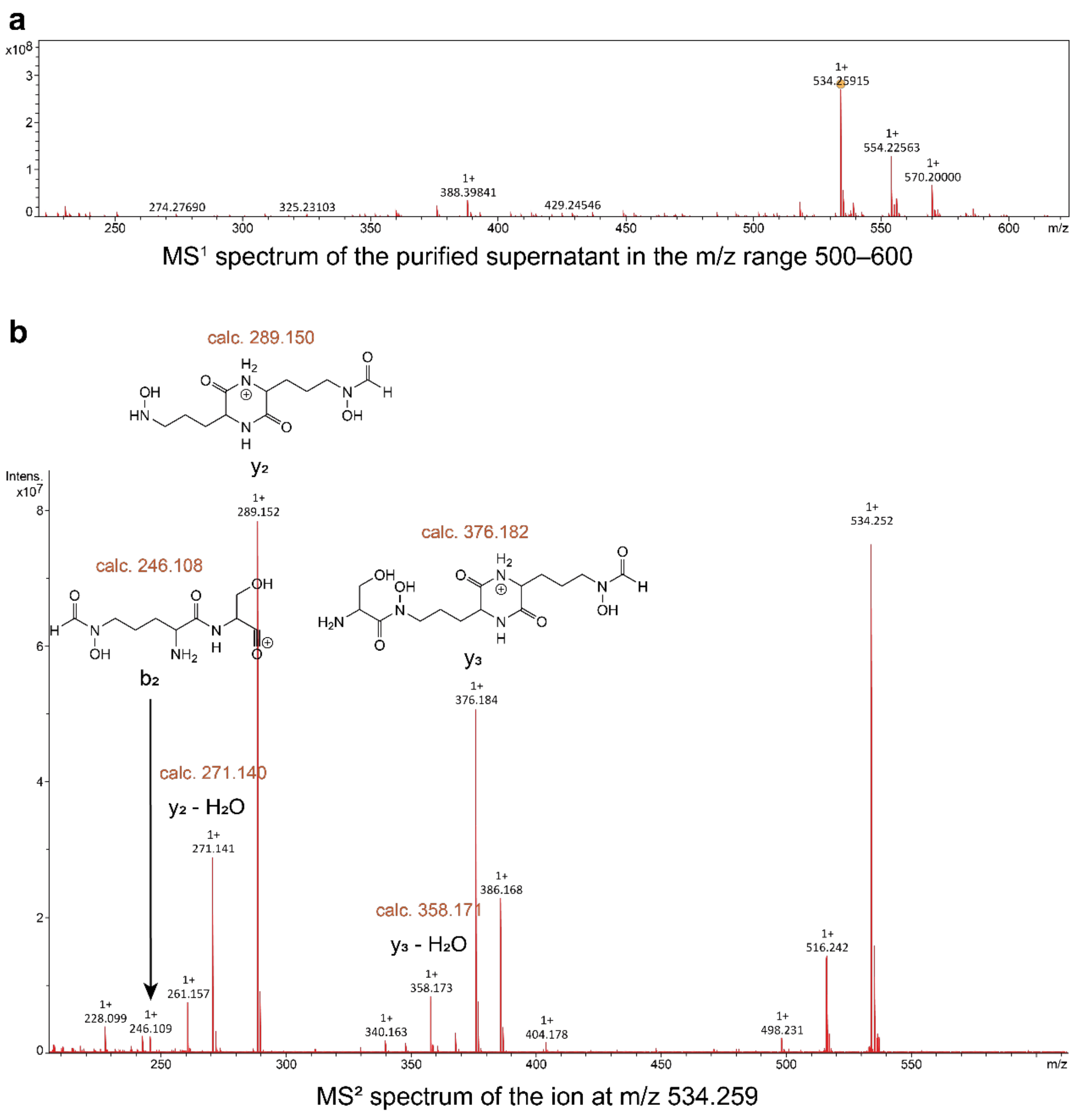


**Fig. S11 | Mass spectrometry confirmation of rhequichelin.** High-resolution MS1 (**a**) and MS/MS fragmentation spectra (**b**) confirming the molecular mass and resolving the structural assembly line of rhequichelin.

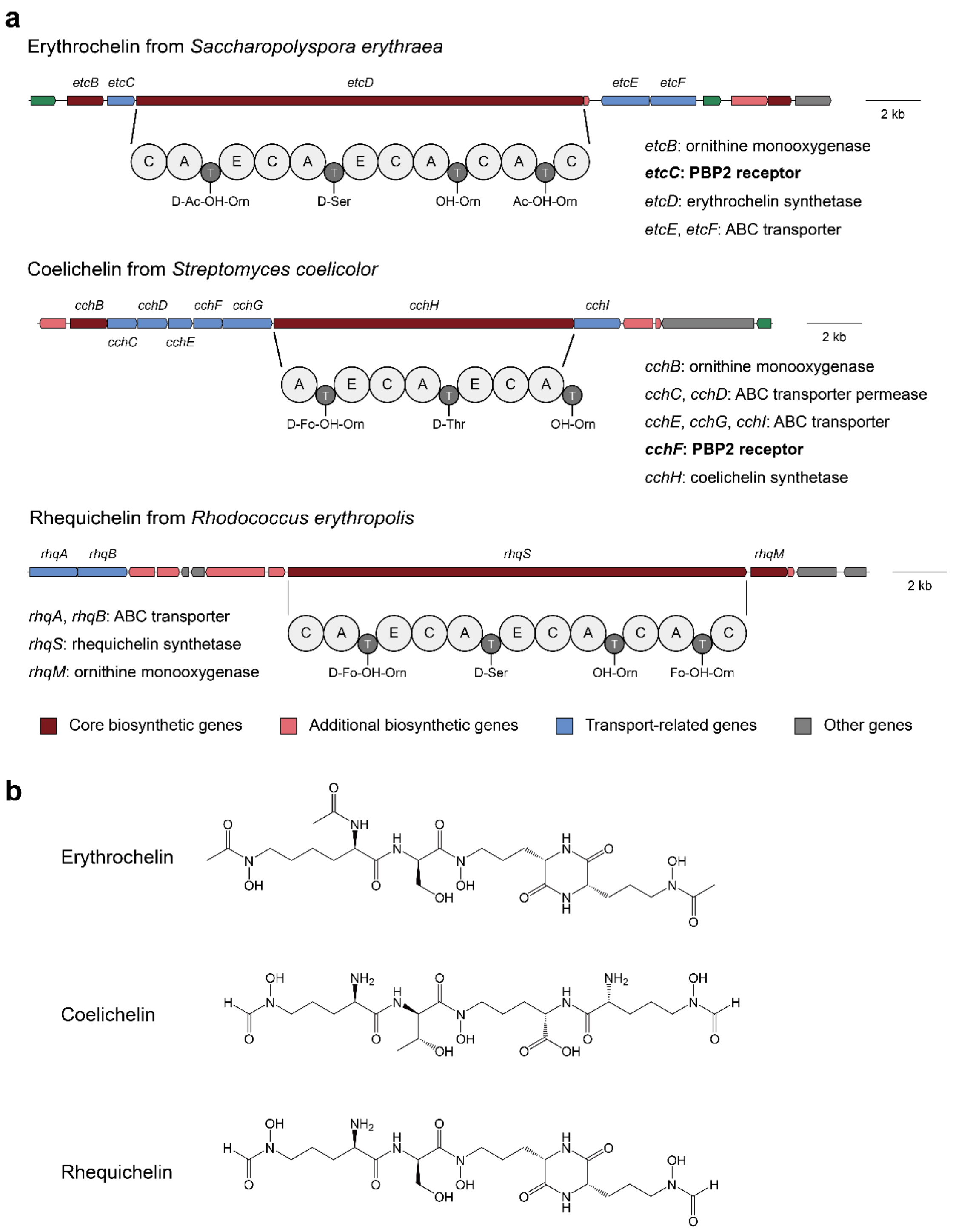


**Fig. S12 | Comparative analysis of the rhequichelin biosynthetic pathway.** Genetic architectures (**a**) and chemical structures (**b**) of the rhequichelin system compared to the structurally related erythrochelin and coelichelin systems, highlighting their shared diketopiperazine (DKP) cyclorelease mechanisms.

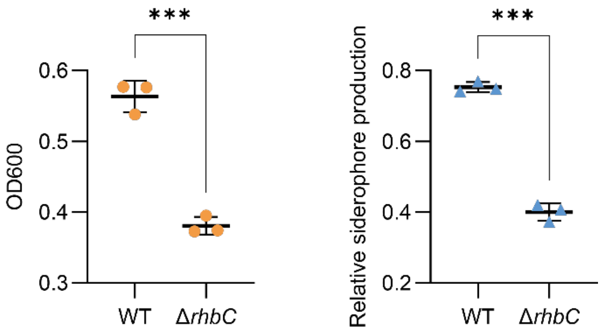


**Fig. S13 | *In vivo* essentiality of the rhequichelin biosynthetic cluster.** Targeted deletion of the *rhbC* cluster (Δ*rhbC*) severely impairs *Rhodococcus* growth under iron limitation (left) and completely abolishes relative siderophore production (right).

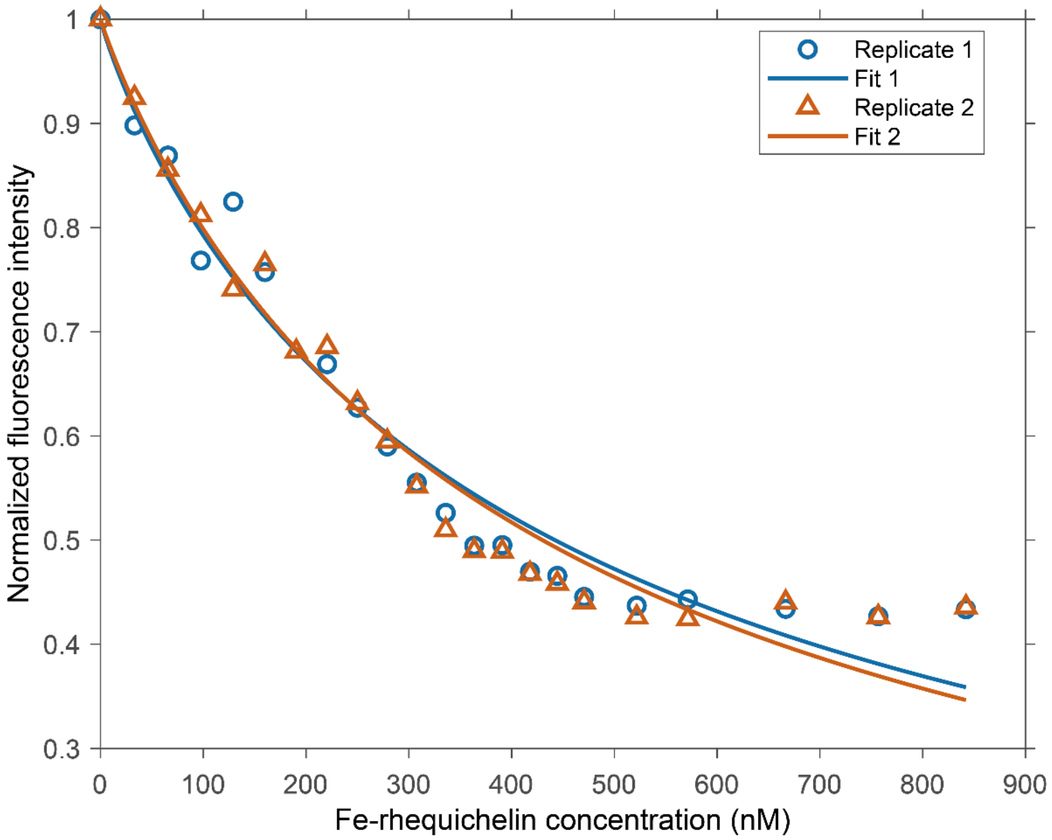


**Fig. S14 | *In vitro* validation of the CIM-predicted RhqR receptor.** Fluorescence quenching assay confirming direct and specific ligand binding between the purified recombinant RhqR receptor and rhequichelin, establishing their functional pairing.

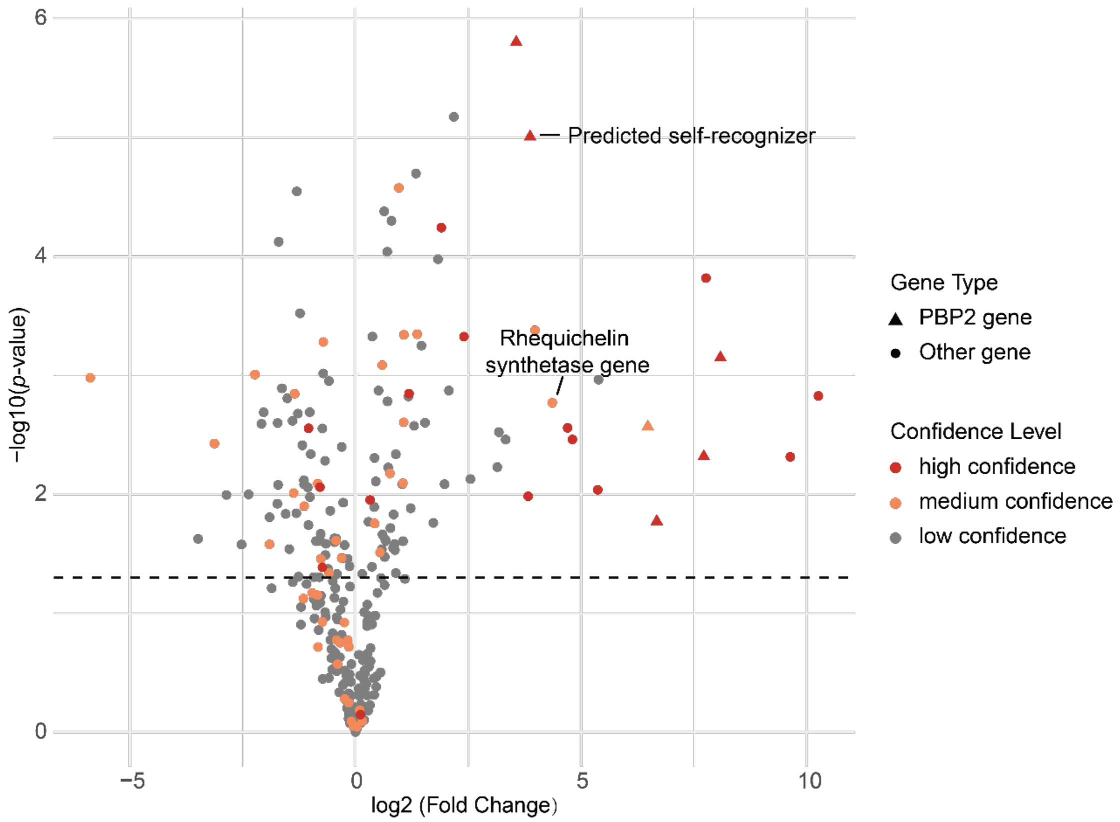


**Fig. S15 | Transcriptomic coordination of the decoupled rhequichelin system.** Volcano plot of the *Rhodococcus* producer strain under iron starvation, revealing the synchronized coordinate upregulation of both the rhequichelin synthetase gene (*rhbC*) and its spatially distant cognate receptor (*rhqR*).
